# Myeloid compartment reprogramming through nanoparticle-delivered resiquimod blocks paracrine growth support and activates phagocytosis to slow tumor progression in endogenous mouse medulloblastoma and diffuse midline glioma models

**DOI:** 10.64898/2026.04.07.714454

**Authors:** Leon McSwain, Kyoungtea Kim, Duhyeong Hwang, Chaemin Lim, Cynthia Winham, Jon Jacques, Hunter Jonus, Elias Rosen, Arpan Pradhan, Sudhir Pai Kasturi, Andrey Tikunov, Serge Yacoub, Dalia Haydar, Alexander V. Kabanov, Jessica Raper, Timothy R. Gershon, Marina Sokolsky-Papkov

## Abstract

In pediatric brain tumors medulloblastoma (MB) and diffuse midline glioma (DMG), tumor-associated myeloid cells (TAMs) support malignant progression by secreting paracrine growth factors and suppressing local immune function. We studied the potential for reversing this cancer-supportive phenotype by stimulating TAM pathogen receptors using ResiPOx, a brain-permeant, polyoxazoline nanoparticle formulation of the TLR7/8 agonist resiquimod. ResiPOx showed blood-brain barrier penetration and anti-tumor efficacy, extending progression-free survival (PFS) in mice with MB and DMG. Integrated cellular and molecular analysis including scRNA-seq showed that ResiPOx expanded TAM populations and reprogrammed TAMs toward anti-tumoral states, blocking paracrine IGF1 signaling and inducing local cytokine signaling and phagocytosis of tumor cells. In rhesus macaques, systemic ResiPOx was well tolerated and induced brain transcriptional patterns that resembled ResiPOx responses in DMG and MB mouse models, indicating effects in non-human primates that highlight translational potential. Our data show that ResiPOx reshapes the brain tumor microenvironment to inhibit tumor growth. As a systemically administered, brain penetrant immunomodulator, ResiPOx is able to reach multifocal and unresectable brain tumors, including MB and DMG.

## INTRODUCTION

Growth-supportive interactions between brain tumors and tumor-associated myeloid cells (TAMs) present opportunities for therapeutic intervention. Myeloid cells, including microglia and bone marrow-derived macrophages (BMDMs), are the most abundant innate immune cells in the central nervous system (CNS). These brain myeloid cells alternate between pro- and anti-inflammatory phenotypes, modulating cytokine and growth factor secretion to coordinate immunosurveillance, response to injury and developmental proliferation ^1–4^. In brain tumors, however, including both medulloblastoma (MB) and high-grade glioma, tumor-associated myeloid cells (TAMs) predominantly show immunosuppressive phenotypes and support tumor growth through paracrine signaling ^5–9^. Here, we examined the potential for pathogen receptor activation to shift TAMs from tumor-supportive to tumor-suppressive states and to produce therapeutic benefits in primary mouse models of pediatric brain tumors, including Sonic Hedgehog (SHH)-driven MB and diffuse midline glioma (DMG).

TAMs play critical roles in SHH MB and DMG. TAMs in SHH MBs secrete IGF1 ligand, which acts as a paracrine growth factor that promotes tumor progression ^9^, by activation IGF receptor (IGF1R) on MB cells, which potentiates SHH signaling ^10, 11^. Reducing TAM populations by treatment with CSF1R inhibitors slows SHH MB progression in mouse models ^12^. However, TAMs in SHH MB are heterogenous ^13^ and while specific TAM populations are associated with tumor growth ^9, 14^, other populations are tumor suppressive ^15, 16^. For example, meningeal macrophages secrete the chemokine CXCL4, which inhibits tumor-promoting CXCL12/CXCR4 signaling, with the effect of slowing SHH MB tumorigenesis ^16^. In DMGs, TAMs secrete growth factors that promote tumor progression ^17, 18^ and adopt immunosuppressive phenotypes ^19, 20^, but retain a capacity for anti-tumor effects through phagocytosis of tumor cells ^21^. Therapies that can enable control of TAM phenotypes may provide new approaches broadly applicable to both MB and DMG.

TAMs express pathogen-associated molecular pattern (PAMP) receptors, including Toll-like receptors 7 and 8 (TLR7/8) ^22, 23^, which mediate dynamic reactions to changes in the tumor microenvironment (TME) ^24, 25^, 20). TLR7/8 activation has received significant attention as a promising immunomodulatory approach in diverse cancers ^26–30^. Resiquimod (R848) is a TLR7/8 agonist that has shown activity with systemic administration in trials for melanoma and other solid malignancies, but has been limited by side effects associated with immune hyper-activation, including cerebral edema ^31–34^. Incorporating resiquimod into a nanoparticle delivery system can mitigate the challenges of poor solubility and both local and systemic toxicities ^30, 35–37^. In brain tumor therapy, resiquimod has shown promise in glioma models when delivered in systemic nanoparticles ^38^ or locally in slow-releasing solid matrix formulation ^39^. These studies showed potent antitumor effects when used to treat mice with implanted syngeneic gliomas, in which the tumor and host derive from different individual animals of the same species.

To advance resiquimod as an improved therapy for multiple types of pediatric brain tumors, we formulated resiquimod in polyoxazoline nanoparticles (ResiPOx), and investigated ResiPOx in mice engineered to develop primary SHH MB or DMG. These models feature endogenous tumors, with intact blood brain barrier (BBB) ^40^ and without potentially immunostimulatory tumor-host mismatch, thus modeling the typically immunologically “cold” phenotype of MBs and DMGs that occur in patients ^20, 41^. Polyoxazoline nanoparticle formulations have shown improved CNS delivery of diverse small molecule agents ^42, 43^ highlighting the potential for ResiPOx to deliver resiquimod across the blood brain barrier directly to the brain tumor TAMs. To investigate this potential, we administered ResiPOx to mice with SHH MB, determined PK and efficacy, and defined pharmacodynamic (PD) effects with cellular and molecular detail. We then compared efficacy in mice with DMG and tolerability and CNS PD in Rhesus macaques, as a model closer in evolution to humans. Our data show that ResiPOx delivers active resiquimod to brain tumors, acts directly on TAMs, synergizes with radiotherapy, and produces similar effects on brain myeloid cells in multiple species, supporting translational potential.

## RESULTS

### ResiPOx enhances delivery across the blood brain barrier and achieves bioactive exposure

We used tritium-labeled resiquimod to compare the PK of ResiPOx or free drug after systemic administration. We generated mice with medulloblastoma by breeding *hGFAP-Cre* and *SmoM2* mouse lines to generate *hGFAP-Cre*/*SmoM2* (*G-Smo*) progeny ^44, 45^. We administered tritium-labeled resiquimod as ResiPOx or free drug to replicate P10 *G-Smo* mice by IP injection. We then harvested the mice at successive time points and analyzed resiquimod concentrations in plasma and tissue samples by scintillation counting, processed with R permutation analysis.

ResiPOx showed higher peak drug concentrations (Cmax) in plasma, forebrain, and tumor (**Fig. 1A**), and showed slower decay of ResiPOx and increased AUC (ΔAUC = 4773.5 p < 0.001 (Plasma), 1523.0 p < 0.0001 (Forebrain), 2265.6 p < 0.0001 (Tumor)) compared to free resiquimod. For orthogonal validation of ResiPOx delivery into the tumor and adjacent brain, we used spectroscopy imaging (MSI) effected by matrix-assisted laser desorption electrospray ionization (IR-MALDESI). This technique applies an infrared laser to frozen tissue sections in an ice matrix and generates concentration maps for all detected metabolites within a present molecular weight range. We mapped the neuronal marker N-acetylaspartic acid (NAA) to distinguish brain from tumor, and mapped creatine to distinguish the edges of the tissue. MSI confirmed detectable accumulation of resiquimod in the tumor and surrounding brain 3 hours after ResiPOx injection, which was no longer detectable by 24 hours (**Fig. 1B**); Together, these data show that ResiPOx enhanced resiquimod PK and successfully delivered the drug into the brain and TME.

**Figure 1.**
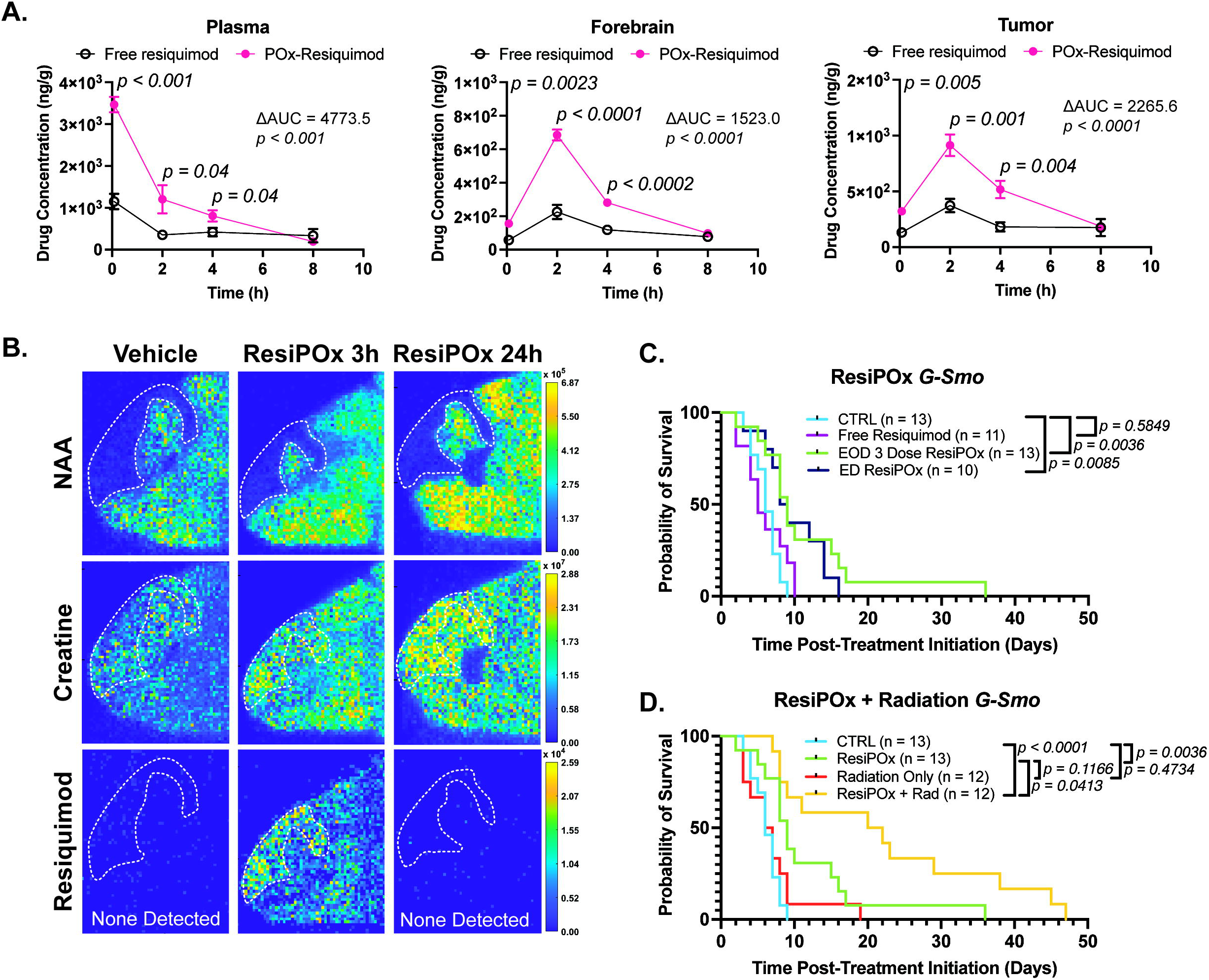
ResiPOx enhances delivery across the blood brain barrier and achieves bioactive exposure in an immunocompetent, genetically engineered mouse model of SHH MB. **(A)** Drug concentration over time in indicated tissues, determined by scintillation counting after administration of tritiated resiquimod, formulated ResiPOx or free drug. Statistical comparisons at individual time points were performed using the Student’s *t*-test and AUCs were compared by timepoint-wise permutation test, n≥4 per timepoint. **(B)** Concentrations of indicated metabolites mapped across the hindbrain and MB (indicated by dotted line) using MSI. Intensity indicates z-score normalized across conditions for each metabolite. **(C)** Kaplan Meyer survival curve demonstrating prolonged survival in *G-Smo* mice treated with either 3 doses of ResiPOx or continuous ResiPOx starting at p10 compared to vehicle or free resiquimod (n≥10, Log-Rank test). **(D)** Kaplan Meyer survival curve demonstrating prolonged survival in *G-Smo* treated with either ResiPOx or Radiation single agent, ResiPOx and radiation combination, or no treatment indicating improved survival in the combination group (n≥12, Log-Rank test).

To determine the potential clinical relevance of ResiPOx-enhanced PK, we compared progression-free survival (PFS) time in *G-Smo* mice treated with ResiPOx or free resiquimod to saline-treated *G-Smo* controls. We randomized *G-Smo* mice to ResiPOx or free resiquimod, administered in three 5 mg/kg doses at P10, P12 and P14. We then followed mouse weight and neurologic status and determined PFS as the time to the humane endpoint of symptomatic tumor progression, detected by sustained weight loss >10% of body weight or onset of neurologic symptoms. ResiPOx-treated mice showed significantly increased PFS compared to either saline or free resiquimod, which were similar (**Fig. 1C**).

ResiPOx was not curative as a single agent in any mice, as all mice showed eventual tumor progression. Because medulloblastoma chemotherapy is typically combined with radiation therapy (RT), we evaluated the efficacy of ResiPOx combined with concurrent RT as a more clinically-relevant approach. Our prior studies showed that *G-Smo* tumors are relatively radiation resistant compared to other genetic mouse MB models ^46^ but show a modest survival benefit with an optimal xRT dose of three fractions of 0.5Gy ^45^. We randomized mice to ResiPOx or saline on P10, P12 and P14, and treated all mice with 0.5 Gy xRT on P11, 13 and 15. We compared survival between these two groups, and to *G-Smo* mice treated with ResiPOx alone shown in **Fig. 1C**. *G-Smo* mice treated with ResiPOx plus RT showed improved PFS compared to either RT alone or ResiPOx alone (**Fig. 1D**), demonstrating synergistic tumor suppression when combining ResiPOx and RT.

### ResiPOx induces changes in TAM and tumor populations

To gain insight into the mechanisms of ResiPOx anti-tumor efficacy, we analyzed ResiPOx effects on TAM populations in *G-Smo* MBs. We randomly assigned *G-Smo* mice to treatment with ResiPOx at P10, P12 and P14 or to no treatment. We then harvested tumors on P15, 24 hours after the last ResiPOx dose, subjected dissociated tumors to flow cytometry and compared treated versus control replicates. Immunostaining for CD45 and CD11b identified discrete TAM subsets in *G-Smo* MBs, including BMDM TAMs, characterized by high CD45 and CD11b expression (**CD45^hi^/CD11B^hi^**) and microglial TAMs with intermediate expression of both markers (**CD45^int^/CD11B^in^**^t^). The microglial TAM compartment comprised two subpopulations, one with relatively higher CD45 expression (**MG^CD45^**) and the other with higher CD11b (**MG^CD11b^**) as shown in **Fig. 2A**. ResiPOx treatment increased the fractions of both BMDM and microglial TAMs, relative to the total cell population (**Fig. 2B**). Within the microglial compartment, the MG^CD11b^ subpopulation increased relative to the total cell population whereas the MG^CD45^ subpopulation showed no significant change (**Fig. 2C**). ResiPOx thus increased TAM populations and showed evidence of change in TAM phenotype.

**Figure 2.**
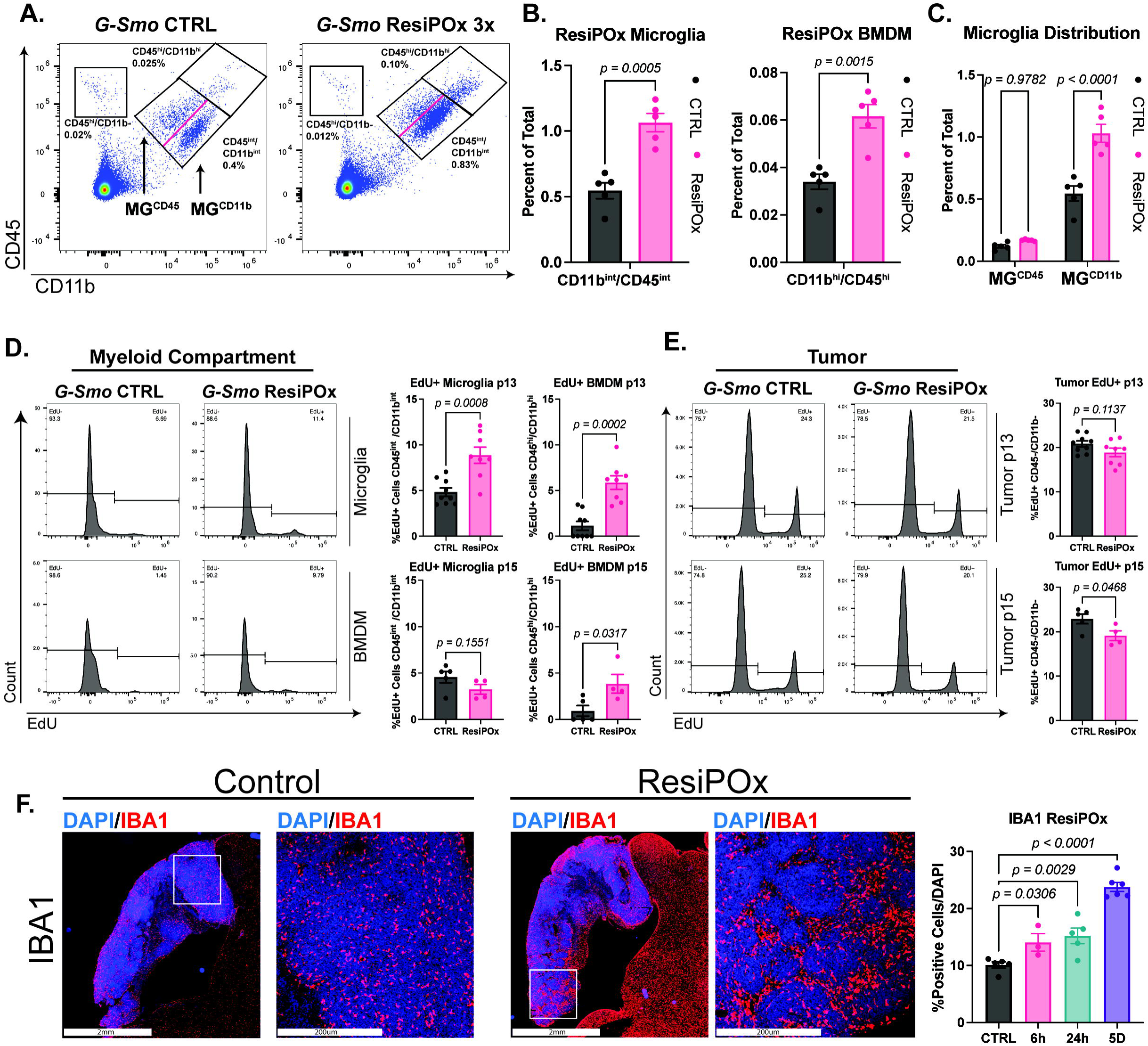
ResiPOx induces changes in TAM and tumor populations. **(A)** Representative scatter plots of MBs from *G-Smo* mice, treated as indicated and then dissociated and subjected to flow cytometry. Antibodies to CD45 and CD11b identify TAMs in the dissociated tumors and distinguish BMDM (CD45^high^/CD11b^high^) and microglial (CD45^int^/CD11b^int^) TAM populations, as well as CD45^+^/CD11b^-^ lymphoid cells, and non-immune populations that were mostly tumor cells (CD45-/CD11b-). **(B,C)** Quantification of each TAM population in replicate mice in each treatment group. (n=5 per group, compared by **(B)** unpaired t-test or **(C)** two-way ANOVA). **(D,E)** Representative EdU studies of **(D)** TAM and **(E)** MB cell populations. ResiPOx treatment in *G-Smo* mice harvested at P13 was administered at P10 and P12, and in P15 *G-Smo* mice harvested at P15 was administered at P10, P12 and P14. Controls were MBs harvested from untreated, age-matched *G-Smo* controls. **(D)** Representative histograms show the fractions of EdU+ cells in the microglial TAMs and BMDM TAMs in MBs of individual P13 G-Smo mice treated as indicated. Graphs quantify EdU labeling microglial TAMs and BMDM TAMs in MBs from replicate mice in each indicated treatment group. (n≥8, *p* determined by Student’s *t*-test). **(E)** Representative histograms as in **(D)** show the fractions of EdU+ tumor cells in individual MBs from G-Smo mice treated as indicated and graphs quantify EdU+ tumor cells in replicates of each treatment group (n≥8, *p* determined by Student’s *t*-test). **(F)** Images show representative IBA1 IHC in hindbrain and MBs from ResiPOx-treated and control *G-Smo* mice (Scale Bar = 2mm or 500um). Graphs show quantification of the IBA1+ fraction of cells in replicate MBs at the indicated times after starting ResiPOx regimen, compared to untreated *G-Smo* MBs (n≥3, *p* determined by one-way ANOVA comparison to NT)

To determine the role of proliferation in the ResiPOx-induced increase in TAMs, we compared *in vivo* EdU uptake in TAMs of ResiPOx-treated and control *G-Smo* mice. We treated one group of *G-Smo* mice with ResiPOx on P10, 12 and P14, administered EdU IP 24 hours later on P15, and another group of *G-Smo* mice with ResiPOx on P10 and 12, administered EdU IP 24 hours later on P13. Tumors from all mice in both groups were harvested 1 hour after EdU injection and compared to age matched untreated *G-Smo* controls, injected IP with EdU IP at P15 or P13. Harvested tumors were dissociated and subjected to flow cytometry. We quantified EdU+ cells in the CD45+/CD11b+ TAM population. This approach also enabled assessment of ResiPOx effects on MB cell proliferation by quantifying EdU+ cells in the CD45-/CD11b- population, which consisted predominantly of MB cells.

Within the TAM population, ResiPOx induced proliferation in both microglial and BMDM subsets at P13, 24 hours after two doses. In BMDMs, this proliferation persisted at P15, 24 hours after 3 doses of ResiPOx (**Fig 2D**). In contrast, in the larger CD45-/CD11b- population that primarily consisted of MB cells, ResiPOx decreased proliferation (**Fig. 2E**).

As an orthogonal approach to quantify TAM populations, we subjected tumors to IHC for the pan-myeloid marker IBA1. We treated sets of replicate *G-Smo* mice with a single IP dose of ResiPOx or saline at P10, harvesting replicates at 6h or 24h after administration. Additionally, we treated sets of replicate *G-Smo* mice with a 3 dose series of ResiPOx or saline at P10, P12 and P14 and harvested tumors 24h later. We then quantified IBA1+ cells in sections of replicate tumors in each condition. We found that ResiPOx increased IBA1+ fractions as early as 6 hours after the first dose and that IBA1+ fractions continued to increase through P16 (**Fig. 2F**). The initial increase at 6h was too rapid to include the effect of proliferation, and most likely reflected increased migration into tumors, while the continued increase included the contribution from increased TAM proliferation.

### ResiPOx induces an acute immune response in MBs and suppresses TAM-to-tumor IGF1 signaling

To identify molecular events induced by ResiPOx, we subjected *G-Smo* MBs from ResiPOx-treated and control mice to transcriptomic analysis. To define acute and chronic transcriptomic changes after ResiPOx without age variation as a potentially confounding variable, we sampled all mice at P14-15.

For acute ResiPOx studies (6h and 24h time points), we administered one dose of ResiPOx to 8 P14 *G-Smo* mice, harvested MBs from 4 replicate mice at 6h or 24h later and compared to 4 *G-Smo* controls injected at P14 with POx-vehicle and harvested 24 hours later. For chronic ResiPOx studies (5D time point), we administered ResiPOx to 4 *G-Smo* mice at P10, P12 and P14 and then harvested MBs at P15 and compared to 4 *G-Smo* controls injected with POx-vehicle at P10, P12 and P14 and harvested at P15. We subjected all MB samples to RNA-seq and compared normalized transcript counts in ResiPOx-treated MBs versus the set of corresponding controls to identify differentially expressed genes (DEGs) with adjusted *p* value < 0.05.

ResiPOx induced maximal transcriptomic changes at 6 hours that decreased with time post-administration. We identified 472 ResiPOx-affected DEGs in MBs at the 6h time point, 313 at 24h, and 131 at 5D (**Fig. 3A–C**). The sets of DEGs at each time point were highly similar, with 240 DEGs commonly affected at 6h and 24h after ResiPOx (p=2e-227 by hypergeometric test) and 55 commonly affected at 24h and 5D (p=2.2e-142 by hypergeometric test) (**Supplementary Fig. 1A,B**). GSEA analysis of DEGs at each time point identified common biologic functions indicating pathogen recognition and innate immune activation (**Fig. 3D and Supplementary Fig 1C,D**). The pattern of DEGs in isolated BMDMs treated for 24h *in vitro* closely resembled the ResiPOx-affected DEGs identified by RNA-seq in *G-Smo* tumors (**Supplementary Fig. 1E,F**), suggesting that the transcriptomic effects induced by ResiPOx in tumors originated from TAMs.

**Figure 3.**
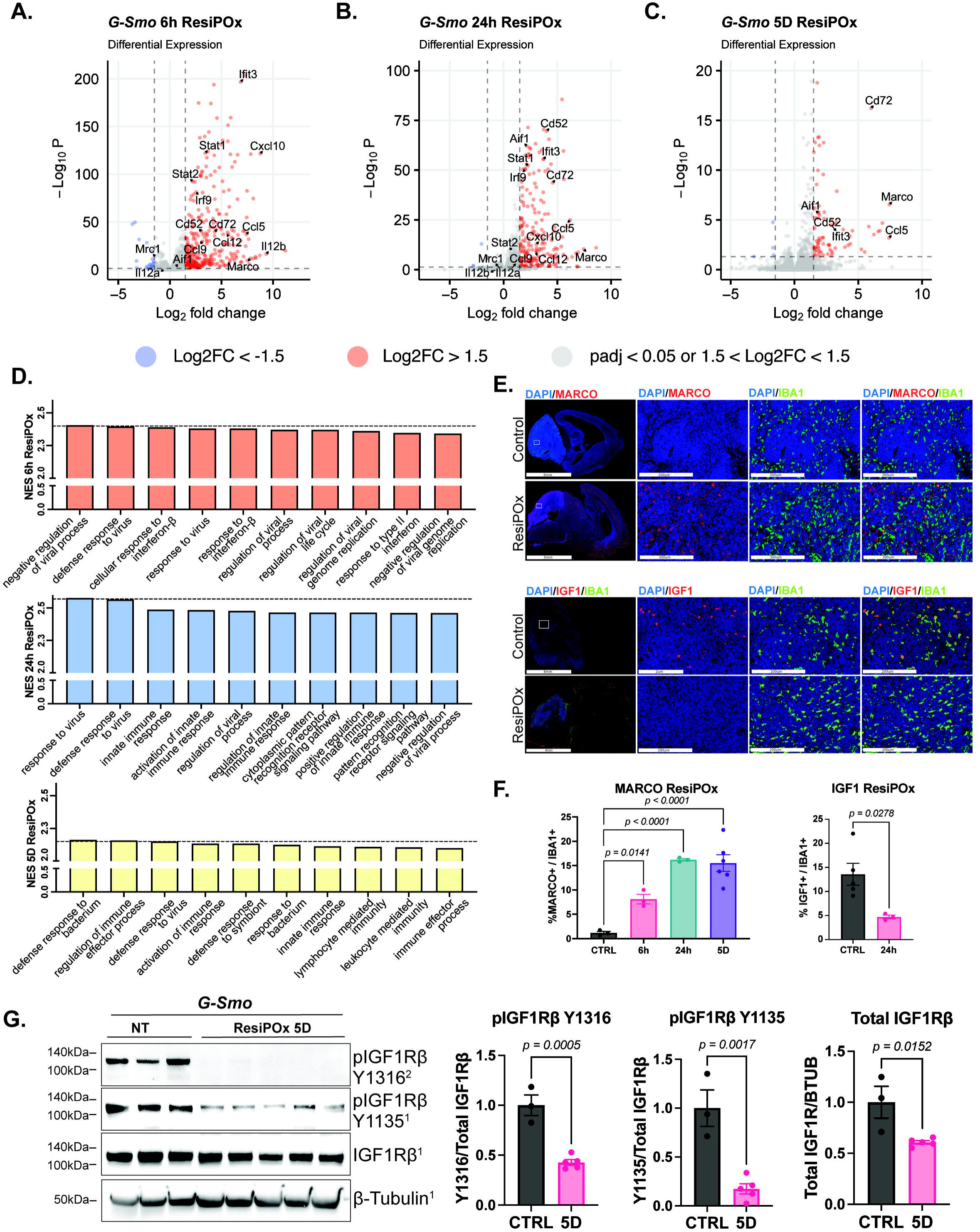
ResiPOx induces a transcriptomic pattern of acute immune response and suppresses IGF1-mediated TAM-tumor signaling. **(A-C)** volcano plots showing DEGs identified by RNA-seq analyses comparing ResiPOx-treated *G-Smo* MBs at the indicated timepoints to age-matched control MBs. Controls for 6h and 24h analyses were untreated P10 *G-Smo* mice. Controls for the 5d time point were untreated P15 *G-Smo* mice. Examples of genes that were differentially induced at multiple time points are labeled, including *Cd72*, *Marco* and *Ccl5*. **(D)** GSEA analysis of ResiPOx-induced DEGs at each time point, showing the 10 pathways with the highest Normalized Enrichment Score (NES), *p < 0.0001 for all pathways*. Pathways common to multiple time points are bolded. **(E)** Images show representative IHC for MARCO and IGF1, both co-localized with the pan-myeloid marker IBA1, in hindbrain and MBs from ResiPOx-treated and control *G-Smo* mice (Scale Bar = 5mm or 300um for MARCO/IBA1 and 4mm or 200um for IGF1/IBA1). Graph show quantification of replicate MBs at indicated time points (n≥3, *p* determined by one-way ANOVA comparison to NT). **(F)** Immunoblot of whole tumor lysates from MBs from untreated P15 *G-Smo* mice compared to MBs from P15 *G-Smo* mice treated with ResiPOx at P10,P12 and P14, showing total and tyrosine phosphorylated IGF1Rβ subunits (pIGF1Rβ1135, pIGF1Rβ1316). Superscript links regions of individual blots to the corresponding loading controls with ^1^ shown in the panel and ^2^ in **Supplementary** Figure 3A. **(G)** Quantification of band intensity in **(F)** normalized to β-Tubulin (n=3 (controls) or 5 (ResiPOx-treated), *p* determined by students t-test).

We used IHC for individual gene products to validate the ResiPOx-induced transcriptomic changes and to identify cell types driving these changes in gene expression. We selected the scavenger receptor MARCO for protein expression studies as it was consistently up-regulated in ResiPOx-treated tumors at 6h, 24h and 5D, was undetectably low in control tumors and showed the highest fold change in the RNA-seq data 5D timepoint. While selected on informatic grounds, MARCO was also of interest as a TAM receptor implicated in tumor control in other cancers ^47^. In sagittal sections containing tumor and adjacent brain, MARCO was undetectable by IHC in control samples, but robustly detected in cells with myeloid morphology. Co-localization with IBA1 confirmed that ResiPOx induced MARCO specifically in myeloid cells, including both TAMs within the MBs and microglia in normal brain regions (**Fig. 3E**). MARCO protein expression began as early as 6 hours after ResiPOx, increased by 24 hours and persisted over 6 days (**Fig. 3F**). These MARCO studies show that TAMs, which comprise a fraction of the tumor population, were the predominant drivers of transcriptomic changes detected in ResiPOx-treated MBs in the bulk RNA-seq dataset.

Along with MARCO, which we selected as a DEG increased with ResiPOx, we studied the protein expression of IGF1, selected as a DEG that decreased with ResiPOx (adj p=0.0042; log2FC=-0.49; Supplementary Data 1). IGF1 is of particular interest as a previous study showed that SHH MBs depend on TAM-derived, IGF1 for paracrine growth factor signaling ^9^. Dual IHC for IGF1 and IBA1 confirmed TAMs as the source of IGF1 in the control *G-Smo* tumors and showed that ResiPOx decreased TAM IGF1 protein expression (**Fig. 3E**). To probe the biologic significance of decreased IGF1, we used immunoblotting to analyzed phosphorylation of the IGF receptor beta subunit (IGF1Rβ) in whole MB lysates. In MB samples from P15 *G-Smo* mice treated with of ResiPOx as in the 5D timepoint, the total IGF1Rβ and the fraction of phosphorylated IGF1Rβ were significantly lower compared to controls (**Fig. 3G and Supplementary. Fig. 2A additional loading controls)**, indicating a decrease in IGF1R activation corresponding with the ResiPOx-suppressed IGF1 ligand. These protein studies show that ResiPOx-induced changes in TAM gene expression could be detected as positive (MARCO) and negative (IGF1) signals in bulk transcriptomic assays, and that produced biologically relevant changes in TAM protein expression and TAM-tumor signaling.

### ResiPOx induces cytokine secretion and IFN**β** signaling

We noted that ResiPOx up-regulated mRNAs coding for cytokines and interferon signaling markers, and that these effects showed variable temporal patterns. We identified 28 cytokines that were up-regulated at 6h and 15 cytokines up-regulated at 24h, of which 13 cytokines were up-regulated at both 6 and 24 hours, and specific cytokines, including *Ccl5*, *Ccl12* and *Ccl19* remained elevated at 24 hours after 3 doses (**Fig 4A**). Similarly, ResiPOx induced multiple interferon response genes (**Fig. 4A**).

**Figure 4.**
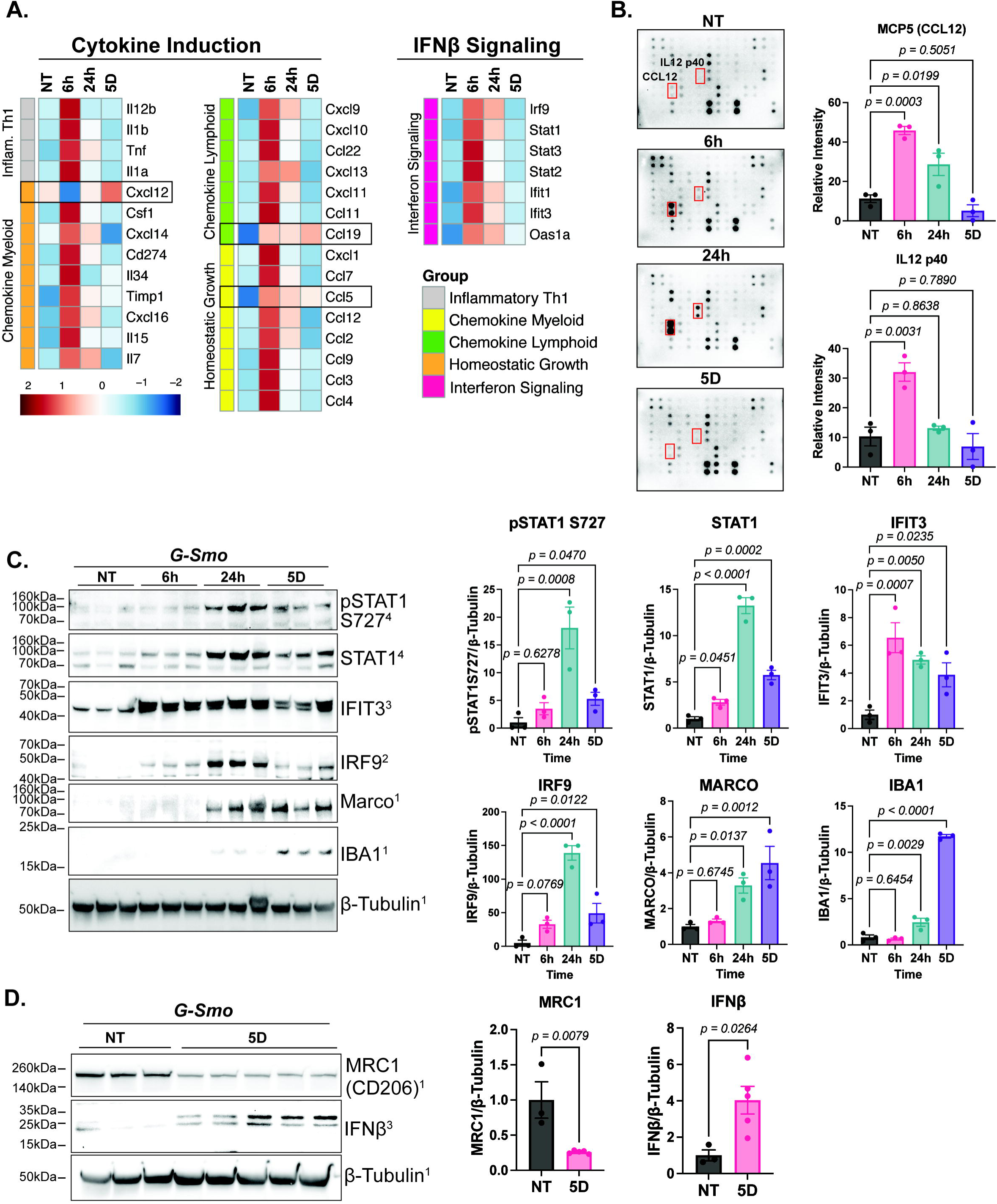
Correlative protein studies show differential expression of proteins encoded by DEGs identified by RNA-seq, including up-regulated cytokines and intracellular interferon effectors and down-regulated MRC1. **(A)** heatmap of RNA-seq data from ResiPOx-treated and control mice from mouse MBs, showing cytokines induced at indicated timepoints after the start of ResiPOx regimen. Boxes indicate induced cytokines that persisted at 5 days. **(B)** Representative cytokine arrays including controls and indicated time points, with quantification of IL12p40 and CCL12 across replicates (n=3 for each condition, *p* determined by one-way ANOVA comparison to NT). Red boxes highlight the positions on the arrays of the two technical replicate locations for IL12p40 and CCL12. **(C)** Immunoblot for indicated proteins on whole tumor lysates from MBs from *G-Smo* mice: NT= untreated P10 *G-Smo* controls, 6h & 24h= *G-Smo* mice harvested 6 or 24 hours after ResiPOx injection at P10, 5d= *G-Smo* mice harvested 24 hours after ResiPOx injection at P10, P12, P14. **(D)** Quantification of band intensity in **(C)**, normalized to β-Tubulin (n=3 for each condition, *p* determined by one-way ANOVA comparison to NT). **(E)** Immunoblot for indicated proteins on whole tumor lysates from MBs from *G-Smo* mice: NT= untreated P15 *G-Smo* controls, 5d= *G-Smo* mice harvested 24 hours after ResiPOx injection at P10, P12, P14. **(F)** Quantification of band intensity in **(E)**, normalized to β-Tubulin (n=3 (controls) or 5 (ResiPOx-treated), *p* determined by students t-test). Superscripts links regions of individual blots to the corresponding loading controls with ^1^ shown in the panel additional controls in **Supplementary** Figure 3D.

Protein studies using cytokine arrays and immunoblot provided orthogonal validation of the RNA-seq data. We compared MB lysates from P10 *G-Smo* mice treated with ResiPOx and harvested at 6h or 24h and P15 *G-Smo* mice treated with ResiPOx at P10, P12 P14 versus a control set of untreated P11 *G-Smo* control mice. A cytokine array that included proteins corresponding to 22 of 28 cytokine DEGs, detected IL12p40 and CCL12 (MCP5) as induced by ResiPOx with maximal signal at 6 hours and waning over time, matching the temporal pattern of *Il12b* and *Ccl12* mRNA (**Fig. 4B, Supplementary Fig. 2B, additional replicates**). On a different array that included an overlapping set of 14 of the 28 cytokine DEGs, CCL9 and CXCL10 were similarly increased (**Supplementary. Fig. 3C**). These protein data were consistent with prior studies showing resiquimod up-regulated IL12p40 ^38, 48^ and confirm RNA-seq data showing that ResiPOx induced specific and dynamic cytokine expression within MBs, consistent with immune engagement.

Immunoblot studies confirmed that ResiPox up-regulated protein expression of multiple interferon signaling proteins, including STAT1, IFIT3 and IRF9, and also showed that ResiPOx induced STAT1 phosphorylation at serine 727 (**Fig. 4C, Supplementary. Fig. 2D additional loading controls**). IFIT3 was induced at 6h and remained elevated until 5D while other interferon signaling proteins including total STAT1, phosphorylated STAT1 S727, and IRF9 peaked at 24h after the first ResiPOx dose and waned by 5D. IFNβ ligand was also elevated at 5D post ResiPOx, varying inversely with anti-inflammatory TAM marker MRC1 (CD206)^49, 50^; ResiPOx thus induced persistent interferon signaling and concurrently suppressed the anti-inflammatory TAM phenotype (**Fig. 4D, Supplementary. Fig. 2A additional loading controls**).

### TAMs are the predominant driver of the transcriptomic response to ResiPOx

To resolve ResiPOx-induced changes with greater cellular resolution, we subjected *G-Smo* mice treated with ResiPOx to scRNA-seq. We compared MBs from 4 *G-Smo* mice treated with ResiPOx at P10, P12 and P14 versus MBs from 4 age-matched, saline-injected *G-Smo* controls. All tumors were dissociated and processed for split-seq-based scRNA-seq using the Evercode Whole Transcriptome (WT) kit v1 (Parse Biotechnologies). Control replicates showed fewer cells sequenced, compared to ResiPOx-treated replicates and to facilitate comparisons we randomly down-sampled the cells from each replicate to match the numbers of cells in the replicate with the smallest number of cells, in keeping with best practices ^45^. After down-sampling, 3865 cells in each replicate passed filtering and were analyzed.

After filtering to remove doublets, unsupervised clustering defined 18 clusters, including 9 discrete clusters and 9 clusters that mapped to a large, multi-cluster group (**Fig. 5A, Supplementary Fig. 3A, Supplementary Data 2**). Cluster-specific DEGs and representative markers identified the discrete clusters as astrocytes, oligodendrocytes, myeloid cells, GABAergic neurons and other non-tumor cells expected in the CNS (**Fig. 5A**, **Table 1**), and identified the multi-cluster group as MB cells in a range of states that paralleled the developmental trajectory of cerebellar granule neuron progenitors (**Table 1**), as in our prior studies in this model ^42, 45, 46, 51, 52^. We used the scRNA-seq data to identify the cell types that expressed the genes that were most strongly induced by ResiPOx in the bulk RNA-seq data (adjusted p< 0.05 and log2 FC > 2). There were 43 genes meeting these criteria in the bulk RNA-seq dataset, and 36 of these 43 genes were detected in the scRNA-seq dataset. The scRNA-seq data showed that all of these 36 genes that were identified as ResiPOx-induced DEGs in the bulk RNA-seq data were specifically expressed by TME cells. While 30/36 ResiPOx-induced DEGs were specific to the myeloid cluster (**Fig. 5B**), 6/36 DEGs, including MHC and interferon response genes were also expressed by endothelial and choroid plexus cells. TAMs were thus the major drivers of the transcriptomic response to ResiPOx, and vascular-facing cells showed an interferon-driven response. TAMs were the only cells to express TLR7, the resiquimod target (**Fig. 5C**), supporting the interpretation that direct drug effects operated through TAMs, and effects on other cell types proceeded indirectly through changes in TAM-mediated signaling.

**Figure 5.**
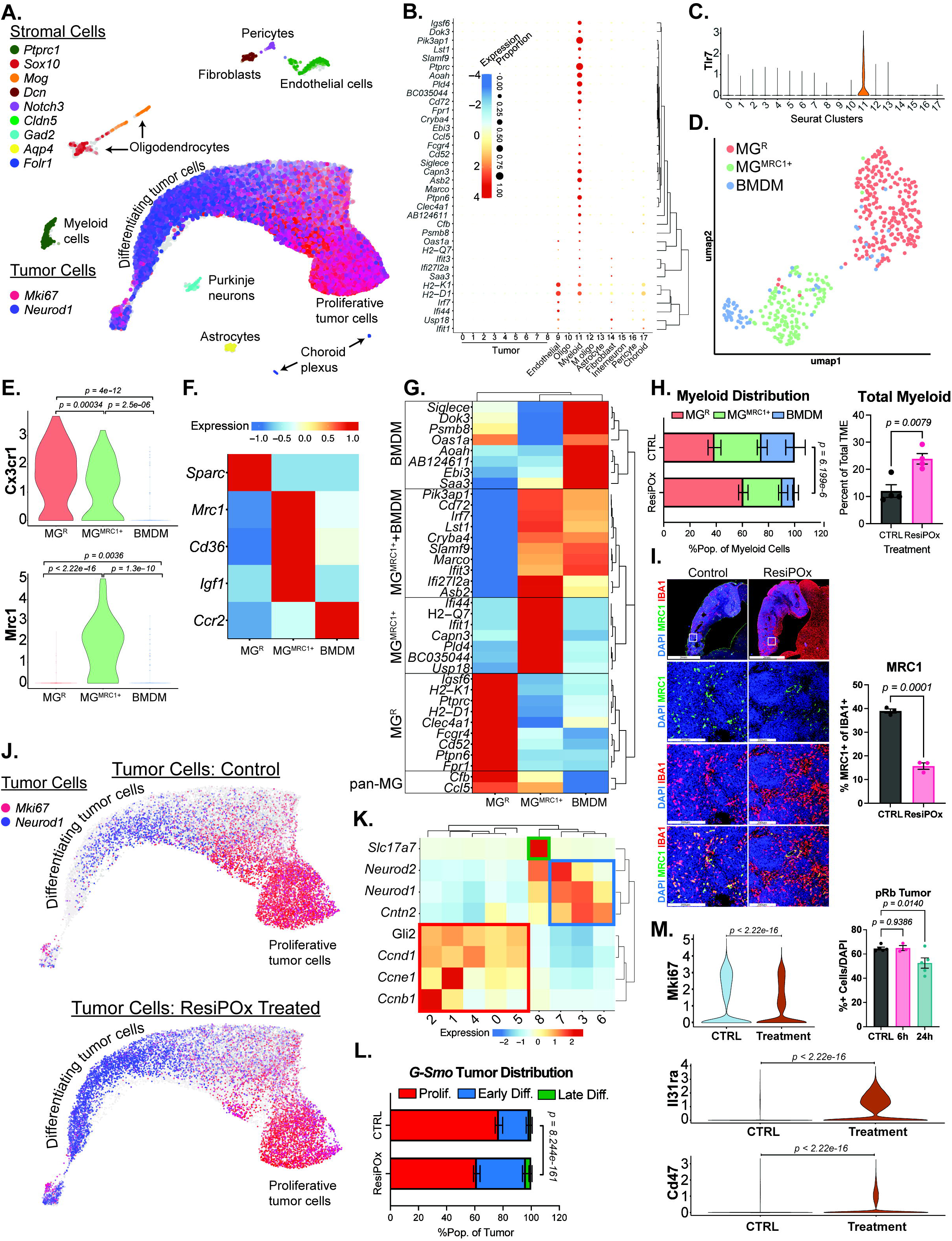
scRNA-seq analysis localizes ResiPOx response TAMs in *G-Smo* MBs, identifies TAM subsets with specific responses and defines indirect effects on tumor cells. **(A)** UMAP showing all cells from treated and control *G-Smo* MBs, with stromal cells color-coded by cell type-specific markers and tumor cells color coded with proliferation marker *Mki67* and differentiation marker *NeuroD1*. Cells are localized according to their proximity to each other in PCA space. **(B)** Bubble plot showing the scaled expression in each tumor cell cluster of DEGs identified in the bulk RNA-seq data as ResiPOx induced (adjusted p<0.05, log2FC >2). Of 43 DEGs in the bulk RNA-seq dataset, 36/43 were detected in the scRNA-seq dataset. Hierarchical clustering of genes grouped the DEGs into 2 major sets, with 30/36 genes expressed specifically by myeloid cells, and 6/36 genes expressed predominantly by endothelial and choroid plexus cells. Oligo= oligodendrocytes and M. Oligo= myelinating oligodendrocytes. **(C)** Violin plot showing expression of TLR7 was specific to the myeloid cluster. **(D)** UMAP showing the cells from the myeloid cluster in the initial PCA, after a second round of PCA and division by cluster analysis into myeloid subclusters, with the cells localized according to proximity in PCA space and color coded by subcluster. **(E)** Violin plots show expression microglial marker Cx3cr1 in MG^R^ and MG^MRC1^ and anti-inflammatory marker Mrc1 limited to MG^MRC1^ (p determined by Wilcoxon rank test). **(F)** Heatmap shows subcluster-specific markers, including *Sparc* in MG^R^, *Igf1* in MG^MRC1^ and *Ccr2* in BMDM. **(G)** Heatmap shows the expression of DEGs as in B, in each myeloid subcluster. Hierarchical clustering of genes divides the DEGs into groups specific to pan-MG, MG^R^, MG^MRC1+^, MG^MRC1+^+BMDM, or BMDM. **(H)** Total myeloid population and distribution of myeloid cells between subclusters, comparing MBs from ResiPOx-treated mice versus to MBs from untreated controls. For distribution comparison *p* = 8.24×10^-161^, determined by Chi square test. For total myeloid comparison, *p* determined by Student’s *t*-test. **(I)** Images show representative IHC co-colocalizing MRC1 and IBA1 in hindbrain and MBs from ResiPOx-treated and control *G-Smo* mice (Scale Bar = 2mm or 200um). Graphs show quantification of the MRC1+ fraction of IBA1+ cells in MBs from replicate *G-Smo* mice in each condition. (n=3 for both groups, *p* determined by one-way ANOVA comparison to NT). **(J)** UMAP as in (A), with only the tumor cells from either ResiPOx-treated or control *G-Smo* mice as indicated. *Mki67* and *NeuroD1* are color-coded. **(K)** Heatmap showing expression of proliferation markers (*Gli2*, *Ccnd1*, *Ccne1, Ccnb1*) and differentiation markers (*Cntn2*, *NeuroD1*, *NeuroD2*, *Slc17a7* (aka *Vglut1*) in each tumor-lineage cluster. Clusters are numbered 0-8 from most populous to least populous. Hierarchical clustering grouped the proliferative clusters and proliferation marker genes (red box) and the early differentiating clusters and early differentiating genes (blue box). Slc17a7, which marks terminal neural differentiation, was limited to cluster 8, termed “late differentiating”, indicated by a green box. **(L)** Distribution of tumor cells between proliferative, early differentiating and late differentiating groups, comparing MBs from ResiPOx-treated mice versus MBs from untreated controls. For distribution comparison, *p* = 6.20×10^-6^, determined by Chi square test. **(M)** Violin plots show expression of indicated genes in tumor cells from ResiPOx-treated versus control MBs (*p* determined by Wilcoxon rank test). Graph shows quantification of pRB IHC in MBs from ResiPOx-treated *G-Smo* mice versus MBs from *G-Smo* controls (*p* determined by one-way ANOVA comparison to NT).

**Table 1.**
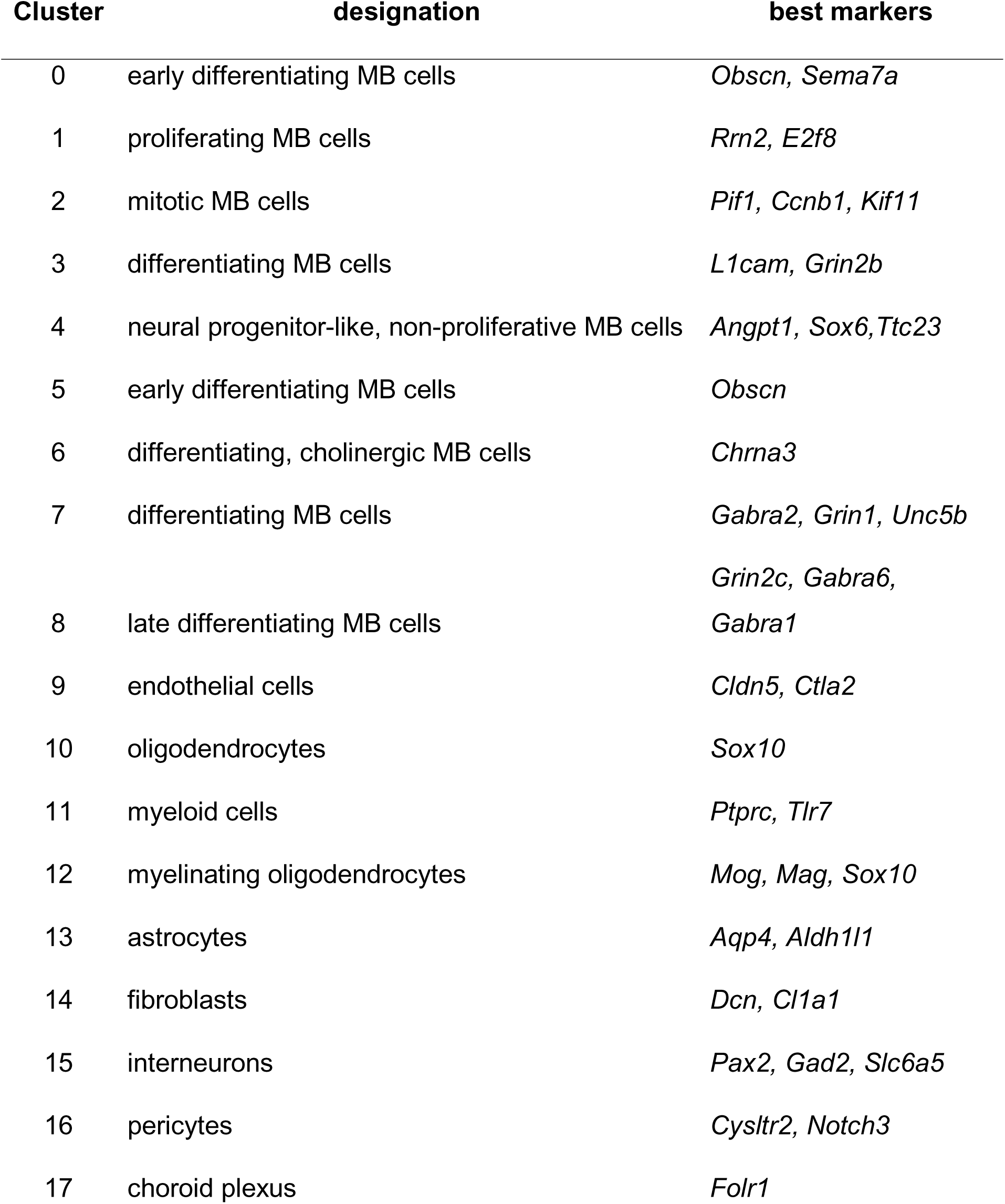
Cell type designations for each cluster in the initial scRNA-seq analysis including both tumor cells and stromal cells. Cluster numbers were assigned in size order, with cluster 0 the most populous.

### ResiPOx alters the proportions of TAMs with specific phenotypes

To investigate the effect of ResiPOx on TAM phenotypes, we isolated the myeloid cluster *in silico* and resolved discrete elements within the TAM population by repeating the PCA and Louvain clustering on the isolated TAMs. This second round of PCA analysis generated a UMAP with three distinct regions, while cluster analysis identified 3 clusters that predominantly correlated with UMAP localization (**Fig. 5D**). Supervised analysis using genes known to mark brain myeloid subsets showed that 2 of the clusters expressed the microglial marker *Cx3cr1* ^53–55^ (**Fig. 5E**), *P2ry12* and *Fcrls* (**Supplementary Fig. 3B,C**) ^13^. One of the two microglial clusters expressed the anti-inflammatory marker *Mrc1*+ ^49, 50^ (*MG^Mrc1+^*; **Fig. 5E**). We further characterized the clusters by determining cluster-specific DEGs (**Table 2, Supplementary Data 3**), which showed that the *Mrc1-* microglial cluster uniquely expressed the ramified microglial marker *Sparc* ^56^ (MG^R^) and MG*^Mrc1+^* expressed anti-inflammatory marker *Cd36* ^57^, supporting an anti-inflammatory role. The *Cx3cr1*- cluster expressed the BMDM markers *Ccr2* ^55^(**Fig. 5F**) and *Cytip* (**Supplementary Fig. 3D**) ^13^. The MG*^Mrc1+^*cluster was the predominant source of *Igf1* (**Fig. 5F**).

**Table 2.** Cell type designations for the myeloid subclustering of the scRNA-seq data. Cluster numbers were assigned in size order, with cluster 0 the most populous.

| Cluster | designation | best markers |
| --- | --- | --- |
| 0 | MG- ramified | *Sparc, Sall1* |
| 1 | MG- MRC1+ | *Mrc1, Cd36* |
| 2 | BMDMs | *Cytip* |

To determine the interaction between ResiPOx and TAM heterogeneity, we determined how each TAM subset contributed to the overall transcriptomic response to ResiPOx, and how ResiPOx affected the relative proportions of TAM subclusters. The MG^R^, and MG*^Mrc1+^* and BMDM subclusters each expressed different subset of the ResiPOx-induced DEGs identified in the bulk RNA-seq (**Fig. 5G**). ResiPOx shifted TAM populations, enriching the MG^R^ (p=6.199 x 10^-6^ Chi square test) at the same time as increasing the overall TAM population relative to all other TME cells (**Fig. 5H**). Across the TAM population, *Mrc1* and *Igf1* were both decreased by ResiPOx (**Supplementary Fig. 3E,F**). Consistent with this change in TAM phenotypes, IHC studies of MRC1 expression, co-localized with pan-myeloid marker IBA1, showed that ResiPOx decreased MRC1+ TAMs (**Fig. 5I**). Together, these scRNA-seq data show that ResiPOx most strongly altered the gene expression in TAMs, induced different, specific patterns of DEGs from different types of TAMs, and shifted TAM phenotypes, enriching MG^R^ TAMs and depleting the *Mrc1*+ TAMs that secrete IGF1.

### scRNA-seq identifies indirect ResiPOx effects on tumor cell gene expression obscured in bulk RNA-seq

While MB cells did not express TLR7 or TLR8 and thus ResiPOx was not expected to alter their gene expression by direct effect, we noted that ResiPOx increased the population of MB cells at the differentiating/differentiated pole in the UMAP (**Fig. 5J**). To investigate the impact of ResiPOx on MB cells in greater detail, we isolated the combined set of 9 clusters of MB cells *in silico*. To facilitate statistical comparisons, we grouped the 9 tumor clusters into three groups according to their position along the developmental trajectory from proliferative, undifferentiated cells to early and late differentiating cells, based on gene expression patterns. Representative markers confirmed the phenotypic characterization of each of these groups, including proliferation markers *Cyclins A1, B1, D1 and E1*, the SHH-driven transcription factor *Gli2*, neural developmental genes *Cntn2*, *Neurod1*, *Neurod2* and the glutamatergic neuron marker *Slc17a7* (aka *Vglut1*; **Fig. 5K**). ResiPOx-treated MBs showed significantly reduced proliferative MB populations and significantly increased early and late differentiating MB populations (**Fig. 5L**), confirming that ResiPOx induced cell cycle exit and differentiation.

We compared gene expression of the combined set of 9 MB clusters in ResiPOx-treated versus control tumors (**Supplementary Data 4**). We noted that ResiPOx increased expression of genes associated with cerebellar granule neuron differentiation, including *Adamts18* ^58^ and *Unc5* ^59^ (**Supplementary Fig. 3G)**. Consistent with inducing a shift in the MB populations toward differentiation, ResiPOx decreased expression of the proliferation marker Mki67 (**Fig. 5M, Supplementary Fig. 3G**). GO Annotation analysis of the set of 253 DEGs with adjusted p<0.05 and log2 FC>1 showed ResiPOx treatment enriched DEGs associated with oxidative phosphorylation (*p*=8.83E-08) including multiple subunits of Cytochrome Oxidase and ATP Synthase and the ADP-ATP exchanger *Slc25a4*^60^. ResiPOx induced DEGs additionally included the immunomodulator *Il31ra* and the phagocytosis inhibitory signal Cd47 (**Fig. 5M**). Analysis of scRNA-seq thus identified ResiPOx-induced changes in MB cells that were obscured by the signal from TME cells in the bulk RNA-seq.

### ResiPOx increased phagocytosis of tumor cells by TAMs

To investigate ResiPOx-induced changes in TAM function, we analyzed effects of ResiPOx on genes related to phagocytosis and antigen presentation. ResiPOx induced genes relevant to both innate and adaptive immune responses including non-opsonic receptors, opsonic Fc receptors, complement cascade components, MHC antigen presentation, and extrinsic apoptosis-inducing ligands and receptors (**Fig. 6A**). Non-opsonic receptors mediate innate recognition of PAMPs and DAMPs. Additionally, opsonic Fc receptors, antigen presentation mechanisms and complement components were also increased, suggesting effects that may link TAMs to humoral responses while upregulation of extrinsic apoptosis pathway components suggests potential activation of TRAIL- and FAS-mediated apoptosis, which may increase phagocytosis of newly killed tumor cells. GSEA confirmed enrichment of transcriptomic patterns consistent with phagocytosis and antigen presentation (**Fig. 6B**). Also consistent with increased phagocytosis, ResiPOx markedly up-regulated expression of BTK **Supplementary Figure 4),** which has been shown to work with TLR activation to promote phagocytosis of cancer cells ^61, 62^.

**Figure 6.**
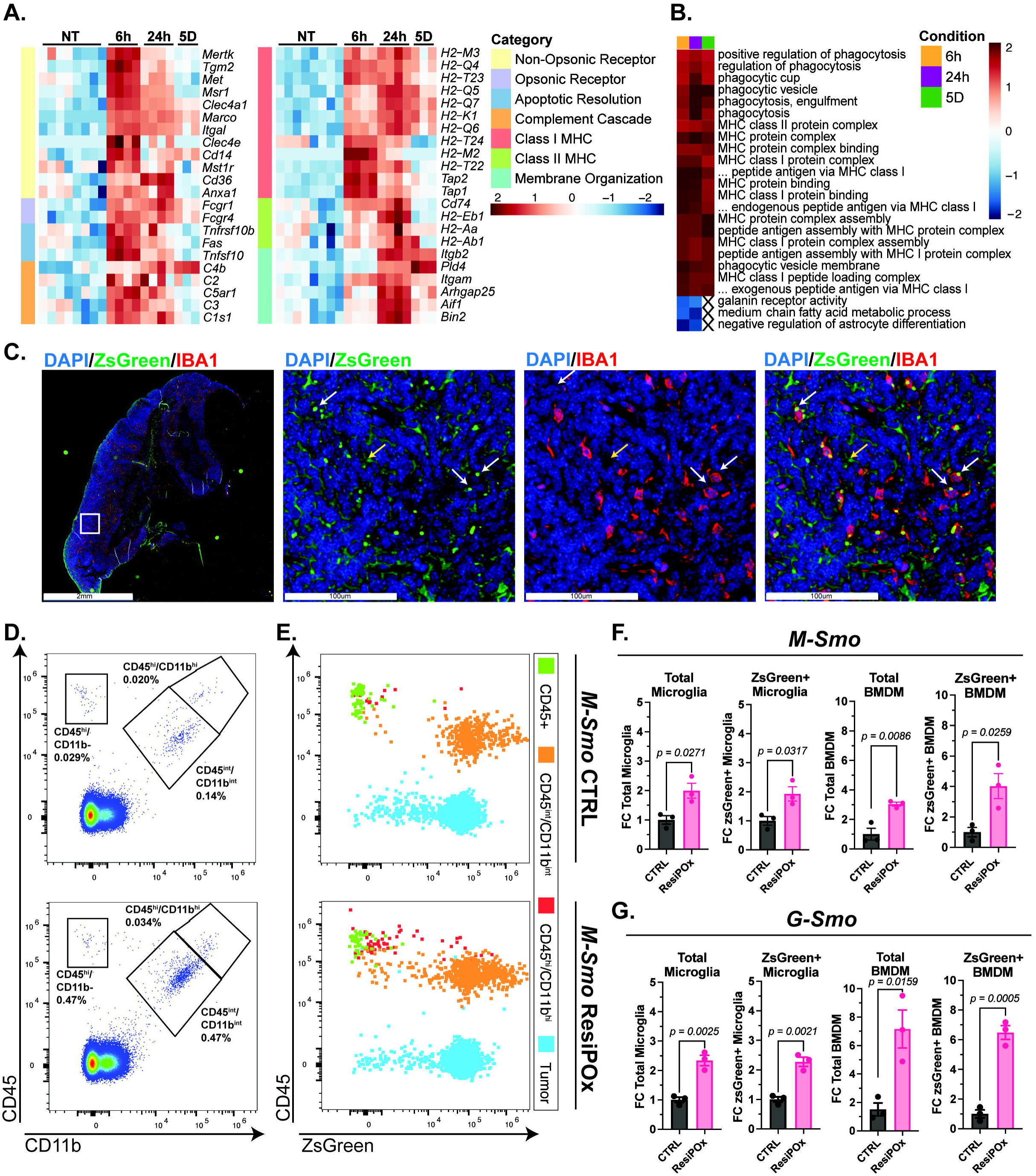
ResiPOx increased phagocytosis of tumor cells by TAMs. **(A)** Heatmap of RNA-seq data from MBs from ResiPOx-treated and control mice, shows in each replicate the expression of genes related to phagocytosis, antigen presentation, apoptotic induction, and membrane organization. MBs from replicate mice are grouped by treatment, as indicated at the top of the heatmap. Only genes with adjusted *p* < 0.05 at least one time point are shown. **(B)** Heatmap of GSEA comparison of MBs from replicate ResiPOx-treated *G-Smo* mice at indicated time point versus MBs from age-matched *G-Smo* controls, showing pathways related to phagocytosis and antigen presentation, with 3 down-regulated pathways selected for comparison. Only pathways with adjusted *p* < 0.05 at least one time point are shown. **(C)** Images show representative *M-Smo^Ai^*^6^ MB, subjected to IHC co-localization of MB-specific ZsGreen and myeloid marker IBA1. ZsGreen is seen in MB cell cytoplasm (yellow arrow) and in intensely staining subcellular structures (white arrows) that co-localize with IBA1, indicating vacuolar localization of tumor-derived zsGreen within TAMs. Scale bars represent 2mm and 100um. **(D)** Representative dot plots of MBs from individual *M-Smo^Ai^*^6^ MBs, treated as indicated and subjected to flow cytometry for CD45 and CD11b, showing the gating used to distinguish BMDM (CD45^high^/CD11b^high^) and microglial (CD45^int^/CD11b^int^) TAM populations, and CD45^+^/CD11b^-^ lymphoid cells. **(E)** Representative dot plot of all Cd45+ cells from ResiPOx-treated or control *M-Smo^Ai^*^6^ MBs, plotting CD45 and ZsGreen, with populations defined by CD45/CD11b color coded. **(F)** Quantification of replicate *M-Smo^Ai^*^6^ tumors treated as indicated and analyzed as in (E), showing the total and zsGreen+ microglial and BMDM TAMs normalized to the total cells per tumor and expressed as fold change (FC) relative to the mean value for controls (n=3, *p* determined by Student’s *t*-test). **(G)** Quantification as in (F) of total and ZsGreen+ microglial and BMDM TAMs in *G-Smo^Ai^*^6^ tumors (n=3, *p* determined by Student’s *t*-test).

To test directly for an effect on phagocytosis, we compared uptake of MB cells by TAM in tumors from mice treated with ResiPOx versus controls. For this purpose, we bred our tumor models with *Ai6* mice, which feature Cre conditional expression of ZsGreen. We bred *Ai6* mice with either *Math1-Cre* mice that express Cre in the cerebellar progenitors that are cells of origin for SHH MB ^63^, or with *Gfap-Cre* mice that express Cre in stem cells that give rise to neurons and glia throughout the brain ^64^. We bred the resulting *Math1-Cre/Ai6* and *Gfap-Cre/Ai6* mice with *SmoM2* mice, to generate pups that developed ZsGreen-expressing MBs. All mice with *Math1-Cre/Ai6/SmoM2* (*M-Smo^Ai^*^6^) and *Gfap-Cre/Ai6/SmoM2* (*G-Smo^Ai^*^6^) genotypes developed MBs as expected. *M-Smo^Ai^*^6^ mice developed MBs with ZsGreen specifically in tumor cells, while *G-Smo ^Ai^*^6^ mice expressed ZsGreen in both MBs and in glial cells throughout the brain. In both genotypes, myeloid cells were outside the lineages that inherited Cre-driven recombination, so that TAMs would only be expected to show ZsGreen fluorescence if they phagocytosed ZsGreen+ cells. IHC for IBA1 and ZsGreen showed ZsGreen+ vacuoles in IBA1+ TAMs (**Fig. 6C**), confirming detection of TAM phagocytosis of MB cells.

To determine the effect of ResiPOx on TAM phagocytosis of MB cells, we treated *M-Smo^Ai^*^6^ and *G-Smo^Ai^*^6^ mice with ResiPOx or sham at P10,P12, P14 and then harvested mice at P15, dissociated MBs and used flow cytometry to quantify ZsGreen+ cells in the CD45^int^/CD11B^int^ microglia and CD45^high^/CD11B^high^ BMDMs (**Fig. 6D**). Representative flow dot plots indicate gating strategy for the myeloid compartment and distribution of ZsGreen+ cells (**Fig. 6 D,E**). Comparison of replicate mice in each genotype and condition showed that ResiPOx increased both total fractions of microglial and BMDM TAMs, and the ZsGreen+ fractions of microglial and BMDM TAMs, relative to the total cell population (**Fig. 6F,G**). ResiPOx thus increased TAM phagocytosis of MB cells, proportionally with the increase in TAM populations.

### ResiPOx shows similar effects in mice with DMG

Considering the common importance of TAMs in MB and in malignant gliomas ^5, 7, 17, 55, 65, 66^, we analyzed ResiPOx efficacy in an immunocompetent DMG model ^67^. We generated mice with DMGs by transducing *Pdgfb*, *Cre*, *H3K27M-Gfp* fusion and *Luciferase* into P2 *Nestin-tva/Tp53^f/f^*mice by intracranial injection of RCAS viruses. We then monitored bioluminescence imaging (BLI) twice weekly to determine the onset of DMG growth. At the first BLI measurement detecting tumor, we initiated 3 IP doses of ResiPOx or POx vehicle, 48 hours apart, as in our MB studies. The glioma model was more slowly progressive than the *G-Smo* MB model, and to address the potential for subjectivity in the identification of glioma symptoms, we randomized mice to ResiPOx or vehicle with the animal handler blinded as to which mice received which treatment. We then continued twice weekly BLI and monitored mice for the onset of symptoms indicating tumor progression, which we defined as the humane endpoint. After 12 mice per group died of progressive tumor, we unblinded the study and analyzed the results.

ResiPOx-treated mice reached the humane endpoint after significantly longer PFS (**Fig. 7A**, *p* = 0.0141). Analysis of BLI measurements over time showed significantly delayed DMG progression in ResiPOx-treated (**Fig. 7B**, *p* < 0.02, determined by a generalized linear model). Individual ResiPOx-treated mice with RCAS-DMG showed transient tumor regression (**Fig. 7C**) that was never seen in controls. Together these data demonstrate ResiPOx efficacy in DMG, comparable to SHH MB.

**Figure 7.**
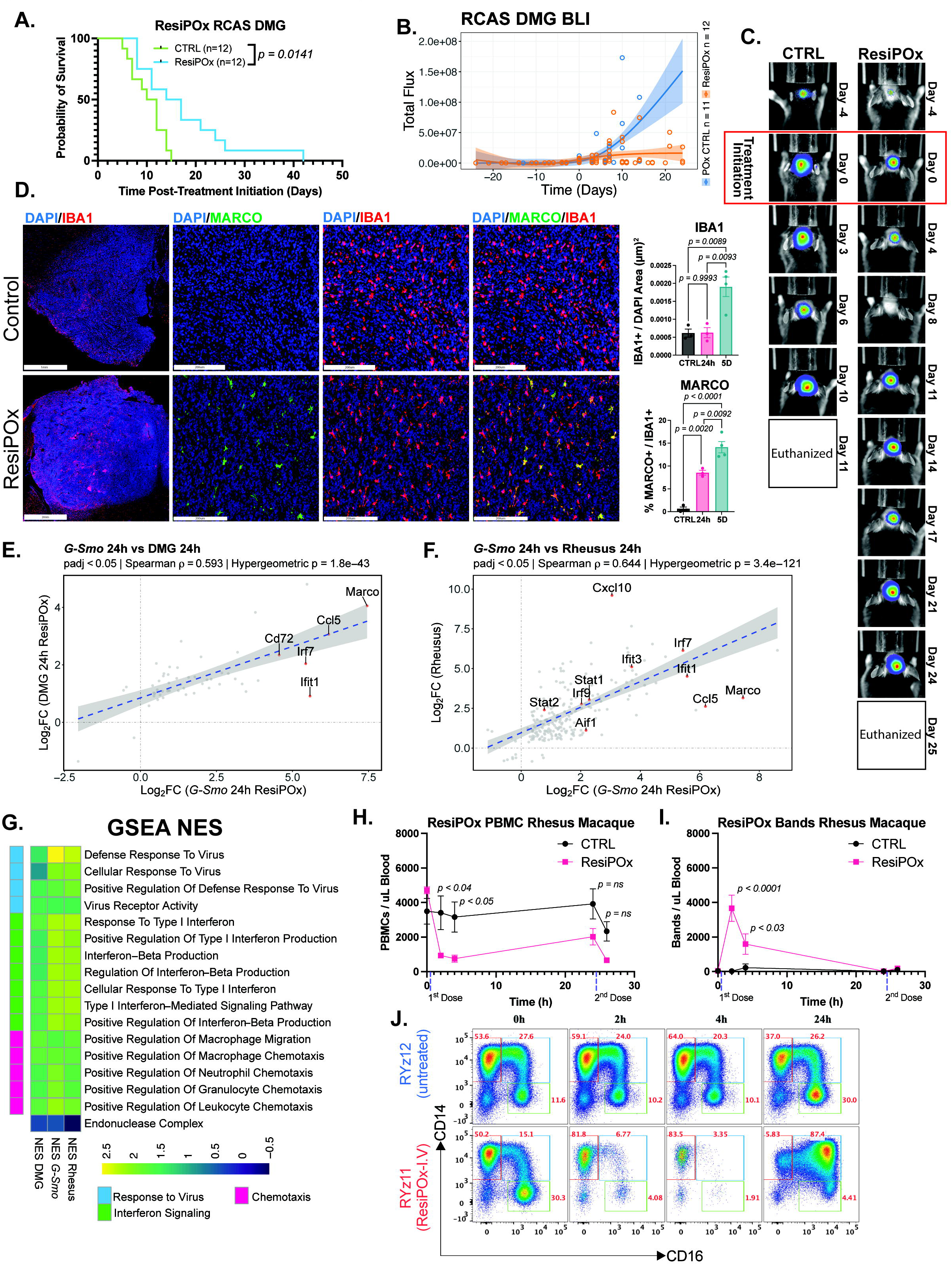
ResiPOx shows efficacy in mice with DMG and similar CNS PD in non-human primates. **(A)** Kaplan Meier curve indicating survival time post-treatment initiation in mice with RCAS-DMG (p = 0.0141, Log-rank Test, n=12 per group). **(B)** BLI signal over time in mice with RCAS DMG subjected to blinded treatment with ResiPOx or POx control formulation. Shaded regions indicate confidence interval and solid line indicates within group fit of natural cubic splines with 3 degrees of freedom calculated using general linearized modeling. Total flux in control mice changed significantly over time compared to ResiPOx treated mice indicating temporal trajectory differences, *p* < 0.02 n ≥ 11 per group. **(C)** Images show representative BLI in RCAS DMG mice treated as indicated. Day 0 indicates time of tumor detection and treatment initiation. **(D)** Images show representative IHC co-localizing MARCO and IBA1 in RCAS DMG mice 24h after 1 dose of ResiPOx or POx control formulation. Graphs show quantification of IBA1+ normalized to DAPI+ area or MARCO+ cells normalized to IBA1+ cells, at indicated time points after the start of the ResiPOx regimen, compared to controls, *p* determined by student’s *t*-test. **(E,F)** Correlation plots of ResiPOx-induced DEGs in *G-Smo* MBs, compared to **(E)** ResiPOx-induced DEGs in RCAS DMGs or **(F)** Rhesus macaque brain samples. In E and F, all samples were harvested 24 hours after administration of ResiPOx (IP in mice, IV in Rhesus macaques) or control (no treatment G-Smo mice, POx vehicle in RCAS DMG mice, saline in Rhesus macaques). Rhesus macaques underwent a second ResiPOx injection 2 hours before harvest. *p* values determined by hypergeometric test. **(G)** heatmap showing NES score for specific ResiPOx-induced pathways commonly activated in *G-Smo* MBs, RCAS DMGs, and Rhesus macaques, determined by GSEA comparison of RNA-seq studies in which ResiPOx treated samples of mouse MB or mouse DMG or macaque brain were harvested 24h after first ResiPOx dose and compared to controls (untreated for G-Smo mice, POx-vehicle-treated for RCAS DMG mice, saline-treated for rhesus macaques). **(H,I)** graphs show quantification of **(H)** PBMCs and **(I)** band neutrophils in peripheral blood in Rhesus macaques after IV administration of ResiPOx or saline (n=3, *p* determined by Student’s *t*-test). **(J)** Dot plots showing CD14 and CD16 divide the monocyte populations from individual treated and control rhesus macaques into classical, non-classical and intermediate subtypes.

As in SHH MB, ResiPOx-treated mice with RCAS DMG showed increased TAM populations and increased MARCO+ TAMs (**Fig. 7D**). We used transcriptomic analysis to compare ResiPOx-induced PD in DMG versus SHH MB. For this study, we generated 4 replicate ResiPOx-treated mice and 3 replicate POx-vehicle controls with RCAS-DMG. We harvested brains from all mice at 24 hours after IP injection and dissected out tumors, guided by GFP expression and processed tumor samples for RNA-seq analysis. We compared tumors from ResiPOx-treated versus control DMGs and identified 193 DEGs (**Supplementary Fig. 7A, Supplementary Data 5**). These ResiPOx-induced DEGs in DMGs showed high concordance with the DEGs up-regulated in MB 24 hours after ResiPOx (*p*=1.7903 x10^-43^ by hypergeometric test), with common induction of *Marco*, *Cd72*, *Ccl5* and *Irf7* (**Fig. 7E**). Together, these results indicate that ResiPOx induces a shared immune-activating program across brain tumors with distinctly different genetics and ontogeny, supporting a common mechanism of action.

### Systemic ResiPOx is well tolerated and shows similar PD in the brain in Rhesus macaques

The efficacy of ResiPOx in multiple mouse brain tumor models supports the clinical potential of ResiPOx for adjuvant brain tumor therapy. To determine ResiPOx safety and PD in the brain in a primate species, we analyzed ResiPOx in adult rhesus macaques at the Emory National Primate Research Center that had pre-existing clinical conditions that warranted a plan for euthanasia, but were able to contribute to this short study prior to euthanasia.

Considering the CNS penetration of ResiPOx in mice, we confirmed safety in non-human primates through dose escalation studies. Using doses of 0.2 mg/kg, 0.4 mg/kg and 0.8 mg/kg in individual animals, we administered 2 IV ResiPOx doses 24 hours apart, monitored animal vital signs and compared complete blood cell counts prior to ResiPOx and 2 hours after the first and second dose. In all 3 animals, all doses were well-tolerated, produced mild, transient temperature elevation. Noting no acute toxicity, we then treated 4 replicate animals, including 2 males and 2 females with 0.8 mg/kg ResiPOx using 2 doses at 24 hours and 2 hours prior to euthanasia. We compared 4 age and sex-matched controls treated with saline.

To analyze CNS effects, we humanely euthanized animals, transcardially perfused with saline and then rapidly harvested brain tissue from animals at 24h after the first ResiPOx or saline injection, which was also 2h after the second injection. Brain samples from hippocampus, parietal cortex and corpus callosum were dissected from similar regions in each animal. We isolated RNA from each brain sample and pooled RNA from each region to generate pooled-region samples from each individual animal. We then sequenced and analyzed the RNA as in our mouse studies, mapping to the Rhesus macaque genome. RNA-seq studies showed that ResiPOx induced significant PD effects in the brain. We identified 1344 DEGs in ResiPOx-treated brain samples (**Supplementary Fig. 6A, Supplementary Data 6**). These DEGs closely matched the ResiPOx induced DEGs in *G-Smo* MBs (*p*=3.4056 x10^-121^ by hypergeometric test) with 216 DEGs commonly induced (**Fig. 7F**). Additionally, comparison of GSEA between *G-Smo* MB, DMG, and Rhesus Macaque indicated common pathways induced by ResiPOx including anti-viral, interferon, and pan-immune chemotaxis (**Fig. 7G)**. Similarly, ResiPOx human macrophages derived from peripheral blood mononuclear cells (PBMCs) and cultured with ResiPOx showed transcriptomic responses that closely resembled mouse BMDMs (**Supplementary Fig. 8, Supplementary Data 7**), indicating a conserved response across species.

To understand the effect of ResiPOx on peripheral blood immune subsets we sampled blood at baseline, and at several time points after each injection. Activation of innate immune cells in peripheral blood with resiquimod has been reported in the context of adjuvant vaccines in rhesus macaques ^68–70^ and we determined the effect of ResiPOx on PBMC frequency and activation when administered I.V at 0.8mg/kg dosing (**Supplementary Fig. 6, Gating Strategy)**. ResiPOx induced changes in circulating myeloid cells, decreasing PBMCs and increasing band neutrophils (**Fig. 7H.I**), consistent with increased passage of myeloid cells from blood to tissues. ResiPOx rapidly increased classical monocytes, up to 83% within 4 hours in comparison to baseline, followed by striking differentiation into intermediate monocytes within 24 hours (**Fig. 7J, Supplementary Fig. 7A)**. Differentiated monocytes displayed striking increases in expression of CD80, CD86 and CD169 in comparison with baseline and isotype controls (**Supplementary Fig. 7B and Supplementary Data 7**). While no substantial changes in frequencies of dendritic cell (DC) subsets were observed compared with saline treated animals, striking increases in the expression of CD80, CD86, CD169 and CD40 were observed on all DC subsets at the 24-hour time point (**Supplementary Fig. 7C,D**). While minimal changes were observed in frequencies and activation of B and T cells, we noted a transient increase in NK cells at 4 hours after ResiPOx (**Supplementary Fig. 7E-G**). These data show appropriate mobilization and activation of blood monocyte and DC subsets that may enable priming of anti-tumor responses in the CNS with ResiPOx treatment. The transcriptomic studies and circulating immune cell studies, taken together, show that ResiPOx crossed the BBB in multiple species, induced a locel gene expression program that was conserved across species, and concurrently promoted immune cell extravasation and tissue infiltration, supporting engagement of shared mechanisms and the potential for similar effects in patients.

## Discussion

These studies shows that ResiPOx delivers resiquimod across the BBB, where it acts on TAMs in MB and DMG, enriching pro-inflammatory TAM phenotypes and inducing cytokine secretion, interferon signaling and phagocytosis of tumor cells, ultimately slowing tumor progression. Immunotherapy for MB and DMG face challenges posed by the BBB, by immunosuppressive myeloid populations that also secrete tumor-supportive growth factors, and by the paucity of immune effectors such as T cells and NK cells ^20, 41, 71^. Our data shows ResiPOx addresses these specific challenges.

ResiPOx showed superior PK and efficacy compared with free resiquimod in *G-Smo* MBs, a genetically-engineered mouse model known to have intact BBB ^40^. ResiPOx induced consistent PD effects within the CNS. In MBs these PD effects included acute induction of cytokine secretion, sustained activation of interferon signaling to vascular cells, increased TAM proliferation and altered TAM phenotypes. ResiPOx induced similar TAM phenotypic changes in DMG, accompanied by comparable anti-tumor efficacy. In rhesus macaques, IV ResiPOx induced transcriptomic changes in normal brain tissue that closely resembled those following IP ResiPOx in mouse brain tumors, indicating consistent BBB penetrance across species and conserved patterns of drug response in normal microglia and in TAMs.

In repolarizing TAMs in *G-Smo* MBs, ResiPOx interrupted IGF1-mediated TAM-to-tumor paracrine growth support. Our scRNA-seq data showed that IGF1 originated specifically from MRC1+ TAMs, consistent with reports localizing IGF1 to MRC1+ myeloid cells outside of the CNS ^72^. ResiPOx decreased MRC1+ TAMs and increased pro-inflammatory TAM populations expressing IL12-p40, matching resiquimod-induced changes in myeloid cell polarization previously described in macrophages *in vitro* ^73^. As microglial TAMs, unlike BMDMs do not circulate outside the CNS, these changes in microglial TAMs indicate local PD effects, in which ResiPOx acted within the CNS compartment to shift TAMs from an immunosuppressive to an activated phenotype that helped suppress tumor growth.

Specific ResiPOx effects on BMDM TAMs indicate systemic effects that were concurrent with local effects in *G-Smo* MBs. The scRNA-seq analysis of *G-Smo* MBs identified discrete patterns of ResiPOx response in BMDM TAMs that differed significantly from microglial TAMs. Moreover, ResiPOx increased proliferation and phagocytosis by BMDM TAMs as well as by microglial TAMs. In our Rhesus macaque studies, which included peripheral blood studies at sequential time points, ResiPOx altered circulating immune populations, markedly reducing PBMCs and transiently increasing band neutrophils. The reduced circulating PBMCs in the Rhesus macaques together with the increased BMDMs in *G-Smo* MBs, suggest migration of circulating immune cells into peripheral organs, as well as BMDM proliferation, may have increased BMDM populations, consistent with prior reports of TLR7-activated myeloid cell migration ^74, 75^. Analysis of activation of blood monocyte and DC subsets highlighted robust activation of CD80, CD86, CD40 and CD169, key nodes in the processes of antigen presentation and engagement of cognate T cells ^76, 77^. These peripheral effects of ResiPOx on circulating immune cells are distinct from, and co-incident with the local ResiPOx effects on microglial TAMs.

A prior study using β-cyclodextrin nanoparticle-formulated resiquimod was the first to show efficacy in glioma ^38^. This work showed that nanoparticle resiquimod administered peripherally to mice with syngeneic, implanted gliomas striking blocked tumor growth through effects on circulating myeloid cells that trafficked to the tumors ^38^. In two recent studies, local administration of matrix-bound resiquimod showed locally-mediated brain tumor suppression ^39, 78^. In our endgenous, immunocompetent glioma model, ResiPOx similarly showed efficacy. In contrast to these other resiquimod delivery approaches, however, ResiPOx acted both peripherally and locally. The local effects of ResiPOx may facilitate additional, concurrent cellular immunotherapies, such as CAR T-cells, which may be stimulated by ResiPOx-induced cytokines including Il12p40 and CXCL10 ^79, 80^. By achieving achieved local delivery of resiquimod to brain tumors without surgery, ResiPOx will be advantageous for patients with tumors not amenable to resection, including recurrent MB and DMG. Additionally, the engagement of the systemic immune system by ResiPOx may contribute to the anti-tumor effect.

While ResiPOx did not detectably increase T cells or NK cells in mouse brain tumors, in a prior study, resiquimod loaded in β-cyclodextrin nanoparticles demonstrated anti-glioma efficacy in mice with global T cell depletion ^38^. In that study, resiquimod increased myeloid cell ROS, suggesting the potential for direct cell killing as an anti-tumor mechanism. The ResiPOx-induced increase in TAM phagocytosis of MB cells may also be a direct anti-tumor mechanism, or may reflect increased tumor cell killing rapidly followed by phagocytosis. Resiquimod therapy thus addresses the lack of immune effectors in brain tumors by recruiting TAMs to this function. Moreover, ResiPOx induced key cytokines that may support exogenous T or NK cell therapies, including CCL9, CXCL10 and CCL12^81, 82^.

In summary, we have developed a nanoparticle platform with distinct advantages for delivery of resiquimod, allowing for systemic administration with safety and tolerability in both mouse models and non-human primates and enabling sustained activation of myeloid cells both systemically and within the TME. Our data are consistent with recent studies showing anti-brain tumor effects of resiquimod delivered systemically in β-cyclodextrin nanoparticles ^38^ or locally in solid matrices ^39, 78^. We now show the efficacy of our agent genetically-engineered primary models of pediatric brain tumors MB and DMG, where endogenous tumors have less potential for tumor-host mismatch. In these models, which are rapidly progressive and highly refractory to therapy, brain tumors arise through a process of tumorigenesis *in situ* and recapitulate intratumoral heterogeneity, TME and treatment-refractory nature of recurrent MB ^46^ and DMG ^67^. Through integrated transcriptomic, protein and cellular analyses in our models, we localized specific features of the resiquimod-induced response to different microglial and BMDM subsets of TAMs and define responses of tumor cells to the disruption caused by ResiPOx. In these models, ResiPOx combined effectively with RT in MB, and ResiPOx produced similar effects in the brain in non-human primates, without apparent toxicity. ResiPOx delivery may be ideal for pediatric brain tumor patients not undergoing resection, including DMGs, which are not resectable, and recurrent MBs, which are typically multifocal and not resected. We conclude that ResiPOx is a promising approach for safe, systemic immunotherapy for pediatric brain tumors that synergizes with RT that is integral to the current standard of treatment.

## Supporting information

Supplementary Data 1. Bulk RNA-seq G-Smo ResiPOx 6 or 24h p 1dose & 5D vs control

Supplementary Data 2. scRNA-seq G-Smo cluster-specific gene lists for combined ResiPOx& control

Supplementary Data 3. scRNA-seq G-Smo TAM subset cluster-specific gene lists for combined ResiPOx& control

Supplementary Data 4. scRNA-seq G-Smo combined tumor clusters ResiPOx v control gene lists

Supplementary Data 5. Bulk RNA-seq DMG ResiPOx 24h p 1dose v control

Supplementary Data 6. Bulk RNA-seq rhesus macaque ResiPOx 2h p 2nd dose vs control

Supplementary Data 7. Bulk RNA-seq human macrophages ResiPOx 24h vs control

## Author Contributions

Y.L., T.D., C.L., T.G.J., L.T., V.V., T.R.G. and B.W.D. conceptualized the manuscript. Y.L., T.D., D.S.M, A.P.T., C.A.P., D.H., C.L., Z.C.B., C.O., U.B., R.C.J.D., S.O., S.K., G.S., B.W., L.T., V.V., L.M., T.R.G. and B.W.D. investigated the findings. Y.L., T.D., C.L., D.S.M., Z.C.B., C.O., S.O., L.T., V.V., B.W., U.B., and L.M. performed the analysis. M.M. and T.H., T.G.J., T.R.G. and G.S. provided the resource. Y.L., T.D., C.L., C.O., L.M. T.R.G., and B.W.D. performed the writing. B.W., M.P., T.R.G. and B.W.D. supervised the findings.

## METHODS

### Materials

We purchased 2-methyl-2-oxazoline (MeOx, 98%), 2-ethyl-2-oxazoline (EtOx, ≥99%), 2-n-butyl-2-oxazoline (BuOx, 99.3%), anhydrous acetonitrile (ACN, for synthesis, 99.8%), and piperidine (≥99.5%) from Millipore Sigma (Burlington, MA, USA). Methyl trifluoromethanesulfonate (MeOTf) (>98.0%) was obtained from Tokyo Chemical Industry (Tokyo, Japan). We acquired resiquimod (Res, >98%) and 3H-Resiquimod (≥96%, 1 mCi mL-1 in EtOH) from MedKoo Biosciences (Durham, NC, USA) and Moravek (Brea, CA, USA), respectively. We procured Ultima gold liquid scintillation cocktail and soluene-350 from Revvity (Waltham, MA, USA). Spectra/Por 3 dialysis membrane tubing (Molecular Weight Cut-Off 3.5 kDa) was purchased from Repligen (Waltham, MA, USA). All antibodies used for IHC, western blots or flow cytometry studies are described in Supplementary Data 8 Materials Table.

### Ethical compliance

This research complies with all relevant ethical regulations. All mouse experiments at UNC were performed with the approval of the UNC IACUC under protocols 19-098 and 21-011. All mouse experiments at Emory University were performed with the approval of the Emory IACUC under protocol 202200148. These protocols all require that mice with brain tumors must be euthanized at the first emergence of symptoms, which is considered the maximum allowable tumor burden. This maximum tumor burden was not exceeded

### Synthesis of Poly(2-oxazoline) block copolymer

Amphiphilic poly(2-oxazoline) block copolymer P(MeOx)_35_-*b*-P(BuOx)_24_-*b*-P(MeOx)_35_ (POx) was synthesized by the living cationic ring-opening polymerization as previously described ^83, 84^. The polymer synthesis was initiated by mixing 1 equivalent (eq) of MeOTf with 35 eq of MeOx in ACN. The polymerization proceeded at 60 °C for 2 days and the degree of polymerization was monitored by nuclear magnetic resonance spectroscopy (NMR). The second and third blocks of the polymer sequentially propagated with 24 eq of BuOx and 35 eq of MeOx in the stated order. To terminate polymerization, we added excess piperidine (> 3 eq) into the reactant and carried out the reaction at 40 °C for 4 hr. Upon completion, the polymer was dissolved in acetone and filtered through a nylon membrane filter (pore size 0.2 μm). The filtrate was dialyzed against deionized water (molecular weight cut-off 3.4 kDa) for a week with semi-daily replacement of water. The dialyzed polymer was lyophilized and stored at 4 °C with desiccant until use. The molecular weight of the polymer (*ca.* 9.2 kDa) was determined by NMR.

### Preparation of ResiPOx

The formulation was prepared by the thin-film method ^85^. Resiquimod and POx were dissolved in 200 proof EtOH to a concentration of 10 mg mL^-1^ and subsequently sonicated at room temperature for 5 min. For ResiPOx preparation, 100 µL of POx solution was mixed with 40 µL of Res solution in a 1.5 mL centrifugation tube (10:4 w/w). The mixture was subjected to drying at 60 °C under nitrogen flow until the solvent completely evaporated. To induce ResiPOx formation, we added 200 µL of saline to the dried thin film. All formulations were freshly prepared from dried thin films immediately prior to administration.

### Determination of ResiPOx pharmacokinetics

To evaluate the PK profile of the formulation in model animals, tritium-labeled ResiPOx (^3^H-ResiPOx) was prepared *via* the thin-film method. (*vide supra*) An EtOH solution of unlabeled Res was mixed with the tritium-labeled Res (^3^H-Res) to yield a total Res concentration of 10 mg mL^-1^ and a radioactivity of 25 µCi mL^-1^ in the ^3^H-ResiPOx. Prior to organ and blood collection at each time point, 100 µL of 1% heparin diluted in saline was subcutaneously administered. Whole blood was collected by cardiac puncture and was centrifuged at 2000 × g for 10 min at 4 °C to separate plasma from whole blood. The plasma fraction from the supernatant was mixed with 1 mL of the 1:1 (v/v) mixture of Soluen-350 and isopropyl alcohol. Harvested organs were completely dissolved in 1 mL of Solene-350 for 3 days. Immediately prior to scintillation counting, 15 mL of Ultima gold liquid scintillation cocktail was added onto each specimen, followed by through vortexting. The level of scintillation from each specimen was determined by Tri-Carb 3110 TR liquid scintillation analyzer (PerkinElmer, Waltham, MA, USA).

### Immunoblotting and Chemokine Array

Whole tumor tissue was isolated into DPBS containing 6g/L glucose and resuspended in 500-1000uL of RIPA + 1x protease/phosphatase inhibitors (RIPA cat CST #9806S, Ptase inhibitors cat CST #5872). Samples were sonicated thoroughly prior to centrifugation and protein quantification using BCA assay. 15-30ug of protein was loaded into each lane of a Tris-Glycine gel and transferred onto 0.45um PVDF immobilon membrane. Membranes were blocked with 5% milk in TBST and probed with respective antibodies in TBST with 5% BSA. For chemokine array (Ray Biotech Cat#AAM-CYT-3-2), 500ug of protein lysate was diluted in array buffer and supplemented with additional protease/phosphatase inhibitors. Lysate was incubated overnight on membranes and washed according to manufacturer’s instructions. Secondary antibodies were then applied overnight followed by wash and development according to manufacturer’s instructions.

### Flow Cytometry

Tumors were first triturated into a single cell suspension using Worthington papain kit (cat#) and subject to RBC lysis. For surface staining, samples were incubated with zombie viability die at 1:500 for 25min, blocked with mouse Fc block for 10 min in FACS buffer, and stained with antibodies to CD45 or CD11b. For surface and intracellular staining, surface proteins were stained as described and subsequently fixed and permed using eBiosciences fix/perm kit (cat# 88-8824-00) according to manufacturer’s instructions and finally intracellularly stained. For cell cycle analysis cells were stained with Ki67 and 7AAD prior to EdU azide reaction using thermo EdU plus flow kit (cat#). All samples were run on Cytek Aurora equipped with 5 laser lines.

The innate immune response in NHP blood was analyzed using flow cytometry as reported earlier (PMID: 32561559). Briefly, PBMCs were isolated from 8 mL CPT tube. The CPT tubes were centrifuged at 2500 rpm for 30 min at room temperature without break. The upper plasma layer was collected and buffy layer containing mononuclear cells were collected in a 15 mL falcon tube. The CPT tubes were washed with 1X DPBS to collect any remaining cells. Finally, the cells were lysed in 2 mL ACK lysis buffer, washed two times with 1X DPBS before resuspending in complete media (RMPI 1640 supplemented with 10% FBS, 1% antibiotic-antimycotic solution). For innate analysis, ∼ 4 millions of cells were first incubated with aqua cell viability staining (Invitrogen, CA) dye at 1:1000 dilution in PBS for 30 minutes at 4 °C. Then cells were washed two times in FACS buffer (1X PBS supplemented with 2% FBS) and following antibodies were used in surface staining for 30 minutes at 4 °C in Brilliant Stain Buffer (BD Biosciences, CA). The BB515 anti-human CD40 (Clone: 5C3, BD Biosciences, CA), PerCP anti-human HLA-DR (Clone: L243, Biolegend, CA), PE anti-human Siglec-1 (CD169) (Clone: 7-239, BD Biosciences, CA), PE/Dazzle 594 anti-human CD1c (Clone: L161, Biolegend, CA), PE-Cy7 anti-human CD123 (Clone: 7G3, BD Biosciences, CA), BV421 anti-human CD80 (Clone: L307.4, BD Biosciences, CA), BV605 anti-human CD14 (Clone: M5E2, BD Biosciences, CA), BV650 anti-human CD86 (Clone: IT2.2, Biolegend, CA), BV786 anti-human CD11c (Clone: SHCL-3, BD Biosciences, CA), Alexa Fluor 647 anti-human Clec9A (CD370) (Clone: 3A4/Clec9A, BD Biosciences, CA), R718 anti-human CD16 (Clone: 3G8, BD Biosciences, CA), BUV395 anti-human CD3 (Clone: SP34-2, BD Biosciences, CA), BUV496 anti-human CD8α (Clone: RPA-T8, BD Biosciences, CA), BUV737 anti-human CD20 (Clone: 2H7, BD Biosciences, CA), BB515 Mouse IgG1, κ isotype control (Clone: X40, BD Biosciences, CA), PE Mouse IgG1, κ isotype control (Clone: MOPC-21, BD Biosciences, CA), BV421 Mouse IgG1, k isotype control (Clone: X40, BD Biosciences, CA), and BV650 Mouse IgG2b, κ isotype control (Clone: MPC-11, Biolegend, CA). Finally, cells were washed and fixed in 150 μL of BD Cytofix™ fixation buffer (BD Biosciences, CA) and acquired in a flow cytometer (BD LSRFortessa, BD Biosciences). The data were analyzed in Flow Jo (v10) software.

### Immunohistochemistry

Brain samples containing MB or DMG were fixed in 4% formaldehyde for 24-48 hours, embedded in paraffin and sections were cut at 4um for MB tissue and 5um for DIPG tissue. MB sections were stained imaged and quantified as previously described ^45, 86^. For DMG, samples were incubated at 60C for 1h followed by de-paraffinization and rehydration and antigen retrieval in sodium citrate buffer in an instant pot at high pressure for 15 minutes. Slide were then photobleached to quench RBCs and background using 3% H2O2 and 20mM NaOH in H2O under activated fluorescent lamps 2x 30min with quench solution change. Samples were blocked with serum and incubated with antibody overnight at 4C followed by HRP-secondary antibody incubation and tyramide amplification reaction. For multiplexing, samples were re-boiled in sodium citrate buffer prior to re-blocking and staining. Slide images were acquired using an Akoya slide scanner with HALO imaging software used for quantification.

### Bulk RNA Sequencing

Tumor samples were harvested in Trizol and flash frozen. After thaw, RNA was purified from Trizol according to manufacturers instructions and treated with DNase prior to paired-end sequencing with 50 million reads. Following sequencing, samples were checked using FastQC, trimmed using TrimGalore, and aligned using STAR genome aligner. Gene reads were normalized and differential genes analysis performed using DEseq in R. For gene set enrichment analysis of DEGs, the fgsea package was used (Korotkevich G et al. 2019). Correlation analysis, including DEGs and GSEA NES values, was performed using Pearson correlation with gene overlap assessed using a hypergeometric test. Principal component analysis of the DMG dataset revealed two samples affected by harvesting-related artifacts. These outliers were excluded prior to subsequent analyses.

### scRNA sequencing

Split-seq based barcoding and library construction was performed using the Parse Fixation and barcoding kits (Cell Fixation Kit v1, Evercode WT (100K) v1) and sequenced on an Illumina Novaseq6000, using an S2 flow cell, as previously described ^45^. The scRNA-seq data were analyzed using Parse software to identify cells by bar code, and to map transcripts to the mouse transcriptome. Data were then analyzed using our previously reported pipeline ^46, 52^.

### MSI

MSI was accomplished using IR-MALDESI, as previously described ^43^. In brief, an IR laser was applied to 10 µm frozen sections of mouse brain with tumor, to desorb adjacent 50µm diameter sampling locations, and an electrospray (50/50 mixture of methanol/water (v/v) with 0.2% formic acid) ionized the desorbed neutral molecules. The resulting ions were sampled into a high resolving power Thermo Fisher Scientific Q Exactive Plus (Bremen, Germany) mass spectrometer for synchronized analysis. The mass spectrometer was operated in positive ion mode from m/z 100 to 400, with resolving power of 140,000FWHM at m/z 200. Resiquimod, creatinine and NAA were identified by m/z through reference to the Human Metabolome Database (hmdb.ca, Canada).

## Acknowledgements

Research reported in this publication was supported in part by the national Institutes of Health (NIH). MS was supported by the National Institute for Neurologic Disease and Stroke (NINDS) (R01NS125073). TRG, AT and LM were supported by the NINDS (R01NS088219, R01NS102627, R01NS106227). AVK was supported by the National Cancer Institute (NCI) (CA264488). Research reported in this publication was supported in part by the Cancer Tissue and Pathology Shared Resource of Winship Cancer Institute of Emory University and NIH/NCI under award number P30CA138292. This research was also supported in part by the National Institutes of Health (NIH) Office of the Director (P51OD011132). The content is solely the responsibility of the authors and does not necessarily represent the official views of the National Institutes of Health. Histopathology/Digital Pathology was also performed by the Pathology Services Core at the University of North Carolina-Chapel Hill. We thank Nicholas Pankow and Edison Floyd in the Pathology Services Core for expert technical assistance with Histopathology and Digital Pathology. The PSC is supported in part by an NCI Center Core Support Grant (P30CA016086) and NC Biotech Impact Innovation Grant (2024-IIG-0015). We thank Kaitlyn Love, Alex van Schoor, Rebecca Richardson and veterinarian, Dr. Fawn Stroud, at the Emory National Primate Research Center for their assistance with data collection on rhesus macaques.

## Author Contributions

L.F.M., D.H., A.V.K., T.R.G. and M.S. conceptualized the manuscript. L.F.M., K.K, D.H., C.L., C.W., J.J., E.R., A.P., S.P.K., A.T., J.R., T.R.G. and M.S. investigated the findings. L.F.M., K.K, D.H., C.L., E.R., A.P., S.P.K., A.T., A.V.K., J.R., T.R.G. and M.S.performed the analysis. A.V.K, J.R., T.R.G. and M.S. provided the resources. L.F.M., S.P.K., A.V.K., J.R., T.R.G. and M.S. performed the writing. A.V.K., J.R., T.R.G. and M.S. supervised the findings.

## LEAD CONTACT

Further information and requests for resources and reagents should be directed to and will be fulfilled by the Lead Contacts, Dr. Timothy Gershon and Dr. Marina Sokolsky-Papkov.

## MATERIALS AVAILABILITY

All unique/stable reagents generated in this study are available from the Lead Contact swith a completed Materials Transfer Agreement.

### Data availability

All transcriptomic data in this study will be deposited in the publicly available GEO database, accession numbers pending.

### Code availability

This paper did not use or create original code. The scRNA-seq analysis was performed using Parse software and publicly available code including Seurat R package version 3.1.1.

### Competing Interests

A.V.K. declares a conflict of interest due to issued and pending patents related to this work and his role as col7lfounder, shareholder, and licensor of technology to DelAQUA Pharmaceuticals, which develops poly(2l7loxazoline)–based drug formulations. M.S. declares a conflict of interest as the spouse of A.V.K. The other authors report no competing interests.

## Notes

### Competing Interest Statement

A.V.K. declares a conflict of interest due to issued and pending patents related to this work and his role as co‑founder, shareholder, and licensor of technology to DelAQUA Pharmaceuticals, which develops poly(2‑oxazoline) based drug formulations. M.S. declares a conflict of interest as the spouse of A.V.K. The other authors report no competing interests.

### Summary of Updates

Manuscript updates authors and adds new data in Supplementary figures.

## References

1. Rusin D, Vahl Becirovic L, Lyszczarz G, Krueger M, Benmamar-Badel A, Vad Mathiesen C, Sigurethardottir Schioth E, Lykke Lambertsen K, Wlodarczyk A. Microglia-Derived Insulin-like Growth Factor 1 Is Critical for Neurodevelopment. Cells. 2024;13(2). Epub 20240118. doi: 10.3390/cells13020184. PubMed PMID: 38247874; PMCID: PMC10813844.

2. Yu D, Jain S, Wangzhou A, Zhu B, Shao W, Coley-O’Rourke EJ, De Florencio S, Kim J, Choi JJ, Paredes MF, Nowakowski TJ, Huang EJ, Piao X. Microglia regulate GABAergic neurogenesis in prenatal human brain through IGF1. Nature. 2025;646(8085):676–86. Epub 20250806. doi: 10.1038/s41586-025-09362-8. PubMed PMID: 40770097; PMCID: PMC12527950.

3. Rodriguez-Gomez JA, Kavanagh E, Engskog-Vlachos P, Engskog MKR, Herrera AJ, Espinosa-Oliva AM, Joseph B, Hajji N, Venero JL, Burguillos MA. Microglia: Agents of the CNS Pro-Inflammatory Response. Cells. 2020;9(7). Epub 20200717. doi: 10.3390/cells9071717. PubMed PMID: 32709045; PMCID: PMC7407646.

4. Colonna M, Butovsky O. Microglia Function in the Central Nervous System During Health and Neurodegeneration. Annu Rev Immunol. 2017;35:441–68. Epub 20170209. doi: 10.1146/annurev-immunol-051116-052358. PubMed PMID: 28226226; PMCID: PMC8167938.

5. Read RD, Tapp ZM, Rajappa P, Hambardzumyan D. Glioblastoma microenvironment-from biology to therapy. Genes Dev. 2024;38(9-10):360–79. Epub 20240625. doi: 10.1101/gad.351427.123. PubMed PMID: 38811170; PMCID: PMC11216181.

6. Persson ML, Douglas AM, Alvaro F, Faridi P, Larsen MR, Alonso MM, Vitanza NA, Dun MD. The intrinsic and microenvironmental features of diffuse midline glioma: Implications for the development of effective immunotherapeutic treatment strategies. Neuro Oncol. 2022;24(9):1408–22. doi: 10.1093/neuonc/noac117. PubMed PMID: 35481923; PMCID: PMC9435509.

7. Chen Z, Giotti B, Kaluzova M, Vallcorba MP, Rawat K, Price G, Herting CJ, Pinero G, Cristea S, Ross JL, Ackley J, Maximov V, Szulzewsky F, Thomason W, Marquez-Ropero M, Angione A, Nichols N, Tsankova NM, Michor F, Shayakhmetov DM, Gutmann DH, Tsankov AM, Hambardzumyan D. A paracrine circuit of IL-1beta/IL-1R1 between myeloid and tumor cells drives genotype-dependent glioblastoma progression. J Clin Invest. 2023;133(22). Epub 20231115. doi: 10.1172/JCI163802. PubMed PMID: 37733448; PMCID: PMC10645395.

8. Zhu C, Mustafa D, Zheng PP, van der Weiden M, Sacchetti A, Brandt M, Chrifi I, Tempel D, Leenen PJM, Duncker DJ, Cheng C, Kros JM. Activation of CECR1 in M2-like TAMs promotes paracrine stimulation-mediated glial tumor progression. Neuro Oncol. 2017;19(5):648–59. doi: 10.1093/neuonc/now251. PubMed PMID: 28453746; PMCID: PMC5464467.

9. Yao M, Ventura PB, Jiang Y, Rodriguez FJ, Wang L, Perry JSA, Yang Y, Wahl K, Crittenden RB, Bennett ML, Qi L, Gong CC, Li XN, Barres BA, Bender TP, Ravichandran KS, Janes KA, Eberhart CG, Zong H. Astrocytic trans-Differentiation Completes a Multicellular Paracrine Feedback Loop Required for Medulloblastoma Tumor Growth. Cell. 2020;180(3):502–20 e19. Epub 2020/01/28. doi: 10.1016/j.cell.2019.12.024. PubMed PMID: 31983537; PMCID: PMC7259679.

10. Fernandez C, Tatard VM, Bertrand N, Dahmane N. Differential modulation of Sonic-hedgehog-induced cerebellar granule cell precursor proliferation by the IGF signaling network. Dev Neurosci. 2010;32(1):59–70. Epub 20100325. doi: 10.1159/000274458. PubMed PMID: 20389077; PMCID: PMC2866582.

11. Rao G, Pedone CA, Del Valle L, Reiss K, Holland EC, Fults DW. Sonic hedgehog and insulin-like growth factor signaling synergize to induce medulloblastoma formation from nestin-expressing neural progenitors in mice. Oncogene. 2004;23(36):6156–62. doi: 10.1038/sj.onc.1207818. PubMed PMID: 15195141.

12. Tan IL, Arifa RDN, Rallapalli H, Kana V, Lao Z, Sanghrajka RM, Sumru Bayin N, Tanne A, Wojcinski A, Korshunov A, Bhardwaj N, Merad M, Turnbull DH, Lafaille JJ, Joyner AL. CSF1R inhibition depletes tumor-associated macrophages and attenuates tumor progression in a mouse sonic Hedgehog-Medulloblastoma model. Oncogene. 2021;40(2):396–407. Epub 20201106. doi: 10.1038/s41388-020-01536-0. PubMed PMID: 33159168; PMCID: PMC7855734.

13. Dang MT, Gonzalez MV, Gaonkar KS, Rathi KS, Young P, Arif S, Zhai L, Alam Z, Devalaraja S, To TKJ, Folkert IW, Raman P, Rokita JL, Martinez D, Taroni JN, Shapiro JA, Greene CS, Savonen C, Mafra F, Hakonarson H, Curran T, Haldar M. Macrophages in SHH subgroup medulloblastoma display dynamic heterogeneity that varies with treatment modality. Cell Reports. 2021;34(13):108917. doi: 10.1016/j.celrep.2021.108917.

14. Lee C, Lee J, Choi SA, Kim SK, Wang KC, Park SH, Kim SH, Lee JY, Phi JH. M1 macrophage recruitment correlates with worse outcome in SHH Medulloblastomas. BMC Cancer. 2018;18(1):535. Epub 2018/05/10. doi: 10.1186/s12885-018-4457-8. PubMed PMID: 29739450; PMCID: PMC5941618.

15. Maximov V, Chen Z, Wei Y, Robinson MH, Herting CJ, Shanmugam NS, Rudneva VA, Goldsmith KC, MacDonald TJ, Northcott PA, Hambardzumyan D, Kenney AM. Tumour-associated macrophages exhibit anti-tumoural properties in Sonic Hedgehog medulloblastoma. Nat Commun. 2019;10(1):2410. Epub 2019/06/05. doi: 10.1038/s41467-019-10458-9. PubMed PMID: 31160587; PMCID: PMC6546707.

16. Pokrajac NT, Tokarew NJA, Gurdita A, Ortin-Martinez A, Wallace VA. Meningeal macrophages inhibit chemokine signaling in pre-tumor cells to suppress mouse medulloblastoma initiation. Dev Cell. 2023;58(20):2015–31 e8. Epub 20230928. doi: 10.1016/j.devcel.2023.08.033. PubMed PMID: 37774709.

17. Ross JL, Chen Z, Herting CJ, Grabovska Y, Szulzewsky F, Puigdelloses M, Monterroza L, Switchenko J, Wadhwani NR, Cimino PJ, Mackay A, Jones C, Read RD, MacDonald TJ, Schniederjan M, Becher OJ, Hambardzumyan D. Platelet-derived growth factor beta is a potent inflammatory driver in paediatric high-grade glioma. Brain. 2021;144(1):53–69. doi: 10.1093/brain/awaa382. PubMed PMID: 33300045; PMCID: PMC7954387.

18. Daginakatte GC, Gutmann DH. Neurofibromatosis-1 (Nf1) heterozygous brain microglia elaborate paracrine factors that promote Nf1-deficient astrocyte and glioma growth. Hum Mol Genet. 2007;16(9):1098–112. Epub 20070330. doi: 10.1093/hmg/ddm059. PubMed PMID: 17400655.

19. Ausejo-Mauleon I, Labiano S, de la Nava D, Laspidea V, Zalacain M, Marrodan L, Garcia-Moure M, Gonzalez-Huarriz M, Hervas-Corpion I, Dhandapani L, Vicent S, Collantes M, Penuelas I, Becher OJ, Filbin MG, Jiang L, Labelle J, de Biagi-Junior CAO, Nazarian J, Laternser S, Phoenix TN, van der Lugt J, Kranendonk M, Hoogendijk R, Mueller S, De Andrea C, Anderson AC, Guruceaga E, Koschmann C, Yadav VN, Gallego Perez-Larraya J, Patino-Garcia A, Pastor F, Alonso MM. TIM-3 blockade in diffuse intrinsic pontine glioma models promotes tumor regression and antitumor immune memory. Cancer Cell. 2023;41(11):1911–26 e8. Epub 20231005. doi: 10.1016/j.ccell.2023.09.001. PubMed PMID: 37802053; PMCID: PMC10644900.

20. Lin GL, Nagaraja S, Filbin MG, Suva ML, Vogel H, Monje M. Non-inflammatory tumor microenvironment of diffuse intrinsic pontine glioma. Acta Neuropathol Commun. 2018;6(1):51. Epub 20180628. doi: 10.1186/s40478-018-0553-x. PubMed PMID: 29954445; PMCID: PMC6022714.

21. Pachocki CJ, Hol EM. Current perspectives on diffuse midline glioma and a different role for the immune microenvironment compared to glioblastoma. J Neuroinflammation. 2022;19(1):276. Epub 20221119. doi: 10.1186/s12974-022-02630-8. PubMed PMID: 36403059; PMCID: PMC9675250.

22. Lu H, Dietsch GN, Matthews MA, Yang Y, Ghanekar S, Inokuma M, Suni M, Maino VC, Henderson KE, Howbert JJ, Disis ML, Hershberg RM. VTX-2337 is a novel TLR8 agonist that activates NK cells and augments ADCC. Clin Cancer Res. 2012;18(2):499-509. Epub 20111129. doi: 10.1158/1078-0432.CCR-11-1625. PubMed PMID: 22128302.

23. Peng J, Tsang JY, Li D, Niu N, Ho DH, Lau KF, Lui VC, Lamb JR, Chen Y, Tam PK. Inhibition of TGF-beta signaling in combination with TLR7 ligation re-programs a tumoricidal phenotype in tumor-associated macrophages. Cancer Lett. 2013;331(2):239–49. Epub 20130111. doi: 10.1016/j.canlet.2013.01.001. PubMed PMID: 23318200.

24. Kawai T, Ikegawa M, Ori D, Akira S. Decoding Toll-like receptors: Recent insights and perspectives in innate immunity. Immunity. 2024;57(4):649–73. doi: 10.1016/j.immuni.2024.03.004. PubMed PMID: 38599164.

25. Kawai T, Akira S. The role of pattern-recognition receptors in innate immunity: update on Toll-like receptors. Nat Immunol. 2010;11(5):373–84. Epub 20100420. doi: 10.1038/ni.1863. PubMed PMID: 20404851.

26. Isvoranu G, Surcel M, Enciu AM, Munteanu AN, Neagu M, Niculae AM, Chiritoiu G, Munteanu CVA, Chiritoiu-Butnaru M. Toll-like Receptor 7/8 Agonists Exert Antitumor Effect in a Mouse Melanoma Model. Medicina (Kaunas). 2026;62(1). Epub 20260109. doi: 10.3390/medicina62010141. PubMed PMID: 41597427; PMCID: PMC12843957.

27. Yazdimamaghani M, Kolupaev OV, Lim C, Hwang D, Laurie SJ, Perou CM, Kabanov AV, Serody JS. Tumor microenvironment immunomodulation by nanoformulated TLR 7/8 agonist and PI3k delta inhibitor enhances therapeutic benefits of radiotherapy. Biomaterials. 2025;312:122750. Epub 20240808. doi: 10.1016/j.biomaterials.2024.122750. PubMed PMID: 39126779; PMCID: PMC11401478.

28. Frega G, Wu Q, Le Naour J, Vacchelli E, Galluzzi L, Kroemer G, Kepp O. Trial Watch: experimental TLR7/TLR8 agonists for oncological indications. Oncoimmunology. 2020;9(1):1796002. Epub 20200721. doi: 10.1080/2162402X.2020.1796002. PubMed PMID: 32934889; PMCID: PMC7466852.

29. Lesniak M, Lipniarska J, Majka P, Kopyt W, Lejman M, Zawitkowska J. The Role of TRL7/8 Agonists in Cancer Therapy, with Special Emphasis on Hematologic Malignancies. Vaccines (Basel). 2023;11(2). Epub 20230128. doi: 10.3390/vaccines11020277. PubMed PMID: 36851155; PMCID: PMC9967151.

30. Vinod N, Hwang D, Azam SH, Van Swearingen AED, Wayne E, Fussell SC, Sokolsky-Papkov M, Pecot CV, Kabanov AV. High-capacity poly(2-oxazoline) formulation of TLR 7/8 agonist extends survival in a chemo-insensitive, metastatic model of lung adenocarcinoma. Sci Adv. 2020;6(25):eaba5542. Epub 20200617. doi: 10.1126/sciadv.aba5542. PubMed PMID: 32596460; PMCID: PMC7299629.

31. Bhagchandani SH, Vohidov F, Milling LE, Tong EY, Brown CM, Ramseier ML, Liu B, Fessenden TB, Nguyen HV-T, Kiel GR, Won L, Langer RS, Spranger S, Shalek AK, Irvine DJ, Johnson JA. Engineering kinetics of TLR7/8 agonist release from bottlebrush prodrugs enables tumor-focused immune stimulation. Science Advances. 2023;9(16):eadg2239. doi: doi:10.1126/sciadv.adg2239.

32. Dudek AZ, Yunis C, Harrison LI, Kumar S, Hawkinson R, Cooley S, Vasilakos JP, Gorski KS, Miller JS. First in human phase I trial of 852A, a novel systemic toll-like receptor 7 agonist, to activate innate immune responses in patients with advanced cancer. Clin Cancer Res. 2007;13(23):7119–25. doi: 10.1158/1078-0432.CCR-07-1443. PubMed PMID: 18056192.

33. Harrison LI, Astry C, Kumar S, Yunis C. Pharmacokinetics of 852A, an imidazoquinoline Toll-like receptor 7-specific agonist, following intravenous, subcutaneous, and oral administrations in humans. J Clin Pharmacol. 2007;47(8):962–9. doi: 10.1177/0091270007303766. PubMed PMID: 17660481.

34. Zahr NM, Zhao Q, Goodcase R, Pfefferbaum A. Systemic Administration of the TLR7/8 Agonist Resiquimod (R848) to Mice Is Associated with Transient, In Vivo-Detectable Brain Swelling. Biology (Basel). 2022;11(2). Epub 20220210. doi: 10.3390/biology11020274. PubMed PMID: 35205140; PMCID: PMC8869423.

35. Shang H, Xia D, Geng R, Wu J, Deng W, Tong Y, Ba X, Zhong Z, He Y, Huang Q, Ye T, Yang X, Jiang K, Peng E, Zhu J, Liu Y, Chen Z, Tang K. Thermosensitive Resiquimod-Loaded Lipid Nanoparticles Promote the Polarization of Tumor-Associated Macrophages to Enhance Bladder Cancer Immunotherapy. ACS Nano. 2025;19(21):19599–621. Epub 20250517. doi: 10.1021/acsnano.4c17444. PubMed PMID: 40380939; PMCID: PMC12139615.

36. Geng R, Shang H, Chen Z, Tang K, Liu Y, Zhu J. ROS-responsive resiquimod prodrug nanoparticles for macrophage reprogramming and enhanced immune checkpoint inhibitor therapy in bladder cancer. Biomaterials. 2026;326:123677. Epub 20250902. doi: 10.1016/j.biomaterials.2025.123677. PubMed PMID: 40907393.

37. Li JX, Shu N, Zhang YJ, Tong QS, Wang L, Zhang JY, Du JZ. Self-Assembled Nanoparticles from the Amphiphilic Prodrug of Resiquimod for Improved Cancer Immunotherapy. ACS Appl Mater Interfaces. 2024;16(20):25665–75. Epub 20240512. doi: 10.1021/acsami.4c01563. PubMed PMID: 38735053.

38. Turco V, Pfleiderer K, Hunger J, Horvat NK, Karimian-Jazi K, Schregel K, Fischer M, Brugnara G, Jahne K, Sturm V, Streibel Y, Nguyen D, Altamura S, Agardy DA, Soni SS, Alsasa A, Bunse T, Schlesner M, Muckenthaler MU, Weissleder R, Wick W, Heiland S, Vollmuth P, Bendszus M, Rodell CB, Breckwoldt MO, Platten M. T cell-independent eradication of experimental glioma by intravenous TLR7/8-agonist-loaded nanoparticles. Nat Commun. 2023;14(1):771. Epub 20230211. doi: 10.1038/s41467-023-36321-6. PubMed PMID: 36774352; PMCID: PMC9922247.

39. Graham-Gurysh EG, Woodring RN, Simpson SR, Mendell SE, Lukesh NR, Pena ES, Moore KM, Ontiveros-Padilla LA, Hendricksen AT, Lopez AM, Williamson GL, Murphy CT, Genito CJ, Hipp KA, Singh G, Zamboni WC, Hingtgen SD, Fecci PE, Bachelder EM, Ainslie KM. Post-resection delivery of a TLR7/8 agonist from a biodegradable scaffold achieves immune-mediated glioblastoma clearance and protection against tumor challenge in mice. Nat Commun. 2025;16(1):8603. Epub 20250929. doi: 10.1038/s41467-025-63692-9. PubMed PMID: 41022761; PMCID: PMC12479796.

40. Phoenix TN, Patmore DM, Boop S, Boulos N, Jacus MO, Patel YT, Roussel MF, Finkelstein D, Goumnerova L, Perreault S, Wadhwa E, Cho YJ, Stewart CF, Gilbertson RJ. Medulloblastoma Genotype Dictates Blood Brain Barrier Phenotype. Cancer Cell. 2016;29(4):508–22. Epub 2016/04/07. doi: 10.1016/j.ccell.2016.03.002. PubMed PMID: 27050100; PMCID: PMC4829447.

41. Eisemann T, Wechsler-Reya RJ. Coming in from the cold: overcoming the hostile immune microenvironment of medulloblastoma. Genes Dev. 2022;36(9-10):514–32. doi: 10.1101/gad.349538.122. PubMed PMID: 35680424; PMCID: PMC9186392.

42. Lim C, Dismuke T, Malawsky D, Ramsey JD, Hwang D, Godfrey VL, Kabanov AV, Gershon TR, Sokolsky-Papkov M. Enhancing CDK4/6 inhibitor therapy for medulloblastoma using nanoparticle delivery and scRNA-seq-guided combination with sapanisertib. Sci Adv. 2022;8(4):eabl5838. Epub 20220126. doi: 10.1126/sciadv.abl5838. PubMed PMID: 35080986; PMCID: PMC8791615.

43. Hwang D, Dismuke T, Tikunov A, Rosen EP, Kagel JR, Ramsey JD, Lim C, Zamboni W, Kabanov AV, Gershon TR, Sokolsky-Papkov Ph DM. Poly(2-oxazoline) nanoparticle delivery enhances the therapeutic potential of vismodegib for medulloblastoma by improving CNS pharmacokinetics and reducing systemic toxicity. Nanomedicine. 2020;32:102345. Epub 2020/12/02. doi: 10.1016/j.nano.2020.102345. PubMed PMID: 33259959.

44. Schuller U, Heine VM, Mao J, Kho AT, Dillon AK, Han YG, Huillard E, Sun T, Ligon AH, Qian Y, Ma Q, Alvarez-Buylla A, McMahon AP, Rowitch DH, Ligon KL. Acquisition of granule neuron precursor identity is a critical determinant of progenitor cell competence to form Shh-induced medulloblastoma. Cancer Cell. 2008;14(2):123–34. doi: 10.1016/j.ccr.2008.07.005. PubMed PMID: 18691547; PMCID: PMC2597270.

45. Li Y, Lim C, Dismuke T, Malawsky DS, Oasa S, Bruce ZC, Offenhauser C, Baumgartner U, D’Souza RCJ, Edwards SL, French JD, Ock LSH, Nair S, Sivakumaran H, Harris L, Tikunov AP, Hwang D, Alicea Pauneto CDM, Maybury M, Hassall T, Wainwright B, Kesari S, Stein G, Piper M, Johns TG, Sokolsky-Papkov M, Terenius L, Vukojevic V, McSwain LF, Gershon TR, Day BW. Suppressing recurrence in Sonic Hedgehog subgroup medulloblastoma using the OLIG2 inhibitor CT-179. Nat Commun. 2025;16(1):1091. Epub 20250204. doi: 10.1038/s41467-024-54861-3. PubMed PMID: 39904981; PMCID: PMC11794477.

46. Malawsky DS, Weir SJ, Ocasio JK, Babcock B, Dismuke T, Cleveland AH, Donson AM, Vibhakar R, Wilhelmsen K, Gershon TR. Cryptic developmental events determine medulloblastoma radiosensitivity and cellular heterogeneity without altering transcriptomic profile. Commun Biol. 2021;4(1):616. Epub 2021/05/23. doi: 10.1038/s42003-021-02099-w. PubMed PMID: 34021242; PMCID: PMC8139976.

47. Xing Q, Feng Y, Sun H, Yang S, Sun T, Guo X, Ji F, Wu B, Zhou D. Scavenger receptor MARCO contributes to macrophage phagocytosis and clearance of tumor cells. Exp Cell Res. 2021;408(2):112862. Epub 20211006. doi: 10.1016/j.yexcr.2021.112862. PubMed PMID: 34626585.

48. Heil F, Hemmi H, Hochrein H, Ampenberger F, Kirschning C, Akira S, Lipford G, Wagner H, Bauer S. Species-specific recognition of single-stranded RNA via toll-like receptor 7 and 8. Science. 2004;303(5663):1526–9. Epub 20040219. doi: 10.1126/science.1093620. PubMed PMID: 14976262.

49. Restagno D, Saraswat M, Aziz PV, Smith K, Roberts AJ, Hintze J, Marth JD. Mrc1 (MMR, CD206) controls the blood proteome in reducing inflammation, age-associated organ dysfunction and mortality in sepsis. Nature Communications. 2025;16(1):6267. doi: 10.1038/s41467-025-61346-4.

50. Stein M, Keshav S, Harris N, Gordon S. Interleukin 4 potently enhances murine macrophage mannose receptor activity: a marker of alternative immunologic macrophage activation. J Exp Med. 1992;176(1):287–92. doi: 10.1084/jem.176.1.287. PubMed PMID: 1613462; PMCID: PMC2119288.

51. Malawsky DS, Dismuke T, Liu H, Castellino E, Brenman J, Dasgupta B, Tikunov A, Gershon TR. Chronic AMPK inactivation slows SHH medulloblastoma progression by inhibiting mTORC1 signaling and depleting tumor stem cells. iScience. 2023;26(12):108443. Epub 20231114. doi: 10.1016/j.isci.2023.108443. PubMed PMID: 38094249; PMCID: PMC10716552.

52. Ocasio J, Babcock B, Malawsky D, Weir SJ, Loo L, Simon JM, Zylka MJ, Hwang D, Dismuke T, Sokolsky M, Rosen EP, Vibhakar R, Zhang J, Saulnier O, Vladoiu M, El-Hamamy I, Stein LD, Taylor MD, Smith KS, Northcott PA, Colaneri A, Wilhelmsen K, Gershon TR. scRNA-seq in medulloblastoma shows cellular heterogeneity and lineage expansion support resistance to SHH inhibitor therapy. Nat Commun. 2019;10(1):5829. Epub 2019/12/22. doi: 10.1038/s41467-019-13657-6. PubMed PMID: 31863004; PMCID: PMC6925218.

53. Zhao XF, Alam MM, Liao Y, Huang T, Mathur R, Zhu X, Huang Y. Targeting Microglia Using Cx3cr1-Cre Lines: Revisiting the Specificity. eNeuro. 2019;6(4). Epub 20190710. doi: 10.1523/ENEURO.0114-19.2019. PubMed PMID: 31201215; PMCID: PMC6620394.

54. Jung S, Aliberti J, Graemmel P, Sunshine MJ, Kreutzberg GW, Sher A, Littman DR. Analysis of fractalkine receptor CX(3)CR1 function by targeted deletion and green fluorescent protein reporter gene insertion. Mol Cell Biol. 2000;20(11):4106–14. doi: 10.1128/MCB.20.11.4106-4114.2000. PubMed PMID: 10805752; PMCID: PMC85780.

55. Chen Z, Feng X, Herting CJ, Garcia VA, Nie K, Pong WW, Rasmussen R, Dwivedi B, Seby S, Wolf SA, Gutmann DH, Hambardzumyan D. Cellular and Molecular Identity of Tumor-Associated Macrophages in Glioblastoma. Cancer Res. 2017;77(9):2266–78. Epub 2017/02/27. doi: 10.1158/0008-5472.CAN-16-2310. PubMed PMID: 28235764; PMCID: PMC5741820.

56. Lloyd-Burton SM, York EM, Anwar MA, Vincent AJ, Roskams AJ. SPARC regulates microgliosis and functional recovery following cortical ischemia. J Neurosci. 2013;33(10):4468–81. Epub 2013/03/08. doi: 10.1523/JNEUROSCI.3585-12.2013. PubMed PMID: 23467362; PMCID: PMC6704956.

57. Hou X, Qu X, Chen W, Sang X, Ye Y, Wang C, Guo Y, Shi H, Yang C, Zhu K, Zhang Y, Xu H, Lv L, Zhang D, Hou L. CD36 deletion prevents white matter injury by modulating microglia polarization through the Traf5-MAPK signal pathway. J Neuroinflammation. 2024;21(1):148. Epub 20240605. doi: 10.1186/s12974-024-03143-2. PubMed PMID: 38840180; PMCID: PMC11155181.

58. Zhu R, Pan YH, Sun L, Zhang T, Wang C, Ye S, Yang N, Lu T, Wisniewski T, Dang S, Zhang W. ADAMTS18 Deficiency Affects Neuronal Morphogenesis and Reduces the Levels of Depression-like Behaviors in Mice. Neuroscience. 2019;399:53–64. Epub 20181221. doi: 10.1016/j.neuroscience.2018.12.025. PubMed PMID: 30579834; PMCID: PMC8006808.

59. Talbot J, Fombonne J, Torrejon J, Babcock BR, McSwain LF, Rama N, Lospinoso Severini L, Bonerandi E, Marsaud V, Bernardi F, Gharsalli T, Guix C, Ducarouge B, Neururer V, Basili I, Mercier AL, Yu H, Forget A, Indersie E, Leboucher S, Souphron J, Okonechnikov K, Wang W, Kawauchi D, Wainwright BJ, Frappaz D, Varlet P, Dufour C, Beccaria K, Blauwblomme T, Martignetti L, Di Marcotullio L, Puget S, Doz F, Bourdeaut F, Masliah-Planchon J, Gershon TR, Mehlen P, Ayrault O. Sonic hedgehog medulloblastomas are dependent on Netrin-1 for survival. Nat Commun. 2025;16(1):5137. Epub 20250603. doi: 10.1038/s41467-025-59612-6. PubMed PMID: 40461501; PMCID: PMC12134228.

60. Bauer MK, Schubert A, Rocks O, Grimm S. Adenine nucleotide translocase-1, a component of the permeability transition pore, can dominantly induce apoptosis. J Cell Biol. 1999;147(7):1493–502. doi: 10.1083/jcb.147.7.1493. PubMed PMID: 10613907; PMCID: PMC2174250.

61. Feng M, Chen JY, Weissman-Tsukamoto R, Volkmer JP, Ho PY, McKenna KM, Cheshier S, Zhang M, Guo N, Gip P, Mitra SS, Weissman IL. Macrophages eat cancer cells using their own calreticulin as a guide: roles of TLR and Btk. Proc Natl Acad Sci U S A. 2015;112(7):2145–50. Epub 20150202. doi: 10.1073/pnas.1424907112. PubMed PMID: 25646432; PMCID: PMC4343163.

62. Benoit RY, Loshi S, Moore CS. Bruton’s Tyrosine Kinase-Going Beyond the B Cell. Immunology. 2026. Epub 20260226. doi: 10.1111/imm.70126. PubMed PMID: 41744142.

63. Machold R, Fishell G. Math1 is expressed in temporally discrete pools of cerebellar rhombic-lip neural progenitors. Neuron. 2005;48(1):17–24. doi: 10.1016/j.neuron.2005.08.028. PubMed PMID: 16202705.

64. Zhuo L, Theis M, Alvarez-Maya I, Brenner M, Willecke K, Messing A. hGFAP-cre transgenic mice for manipulation of glial and neuronal function in vivo. Genesis. 2001;31(2):85–94. Epub 2001/10/23. PubMed PMID: 11668683.

65. Ross JL, Puigdelloses-Vallcorba M, Pinero G, Soni N, Thomason W, DeSisto J, Angione A, Tsankova NM, Castro MG, Schniederjan M, Wadhwani NR, Raju GP, Morgenstern P, Becher OJ, Green AL, Tsankov AM, Hambardzumyan D. Microglia and monocyte-derived macrophages drive progression of pediatric high-grade gliomas and are transcriptionally shaped by histone mutations. Immunity. 2024;57(11):2669–87 e6. Epub 20241011. doi: 10.1016/j.immuni.2024.09.007. PubMed PMID: 39395421; PMCID: PMC11578068.

66. Bhatia B, Northcott PA, Hambardzumyan D, Govindarajan B, Brat DJ, Arbiser JL, Holland EC, Taylor MD, Kenney AM. Tuberous sclerosis complex suppression in cerebellar development and medulloblastoma: separate regulation of mammalian target of rapamycin activity and p27 Kip1 localization. Cancer Res. 2009;69(18):7224–34. Epub 2009/09/10. doi: 0008-5472.CAN-09-1299 [pii]10.1158/0008-5472.CAN-09-1299. PubMed PMID: 19738049; PMCID: 2745891.

67. Cordero FJ, Huang Z, Grenier C, He X, Hu G, McLendon RE, Murphy SK, Hashizume R, Becher OJ. Histone H3.3K27M Represses p16 to Accelerate Gliomagenesis in a Murine Model of DIPG. Mol Cancer Res. 2017;15(9):1243–54. Epub 20170518. doi: 10.1158/1541-7786.MCR-16-0389. PubMed PMID: 28522693; PMCID: PMC5581686.

68. Kasturi SP, Rasheed MAU, Havenar-Daughton C, Pham M, Legere T, Sher ZJ, Kovalenkov Y, Gumber S, Huang JY, Gottardo R, Fulp W, Sato A, Sawant S, Stanfield-Oakley S, Yates N, LaBranche C, Alam SM, Tomaras G, Ferrari G, Montefiori D, Wrammert J, Villinger F, Tomai M, Vasilakos J, Fox CB, Reed SG, Haynes BF, Crotty S, Ahmed R, Pulendran B. 3M-052, a synthetic TLR-7/8 agonist, induces durable HIV-1 envelope-specific plasma cells and humoral immunity in nonhuman primates. Sci Immunol. 2020;5(48). doi: 10.1126/sciimmunol.abb1025. PubMed PMID: 32561559; PMCID: PMC8109745.

69. Kasturi SP, Kozlowski PA, Nakaya HI, Burger MC, Russo P, Pham M, Kovalenkov Y, Silveira ELV, Havenar-Daughton C, Burton SL, Kilgore KM, Johnson MJ, Nabi R, Legere T, Sher ZJ, Chen X, Amara RR, Hunter E, Bosinger SE, Spearman P, Crotty S, Villinger F, Derdeyn CA, Wrammert J, Pulendran B. Adjuvanting a Simian Immunodeficiency Virus Vaccine with Toll-Like Receptor Ligands Encapsulated in Nanoparticles Induces Persistent Antibody Responses and Enhanced Protection in TRIM5alpha Restrictive Macaques. J Virol. 2017;91(4). Epub 20170131. doi: 10.1128/JVI.01844-16. PubMed PMID: 27928002; PMCID: PMC5286877.

70. Kwissa M, Nakaya HI, Oluoch H, Pulendran B. Distinct TLR adjuvants differentially stimulate systemic and local innate immune responses in nonhuman primates. Blood. 2012;119(9):2044–55. Epub 20120112. doi: 10.1182/blood-2011-10-388579. PubMed PMID: 22246032; PMCID: PMC3311246.

71. Price G, Bouras A, Hambardzumyan D, Hadjipanayis CG. Current knowledge on the immune microenvironment and emerging immunotherapies in diffuse midline glioma. EBioMedicine. 2021;69:103453. Epub 20210619. doi: 10.1016/j.ebiom.2021.103453. PubMed PMID: 34157482; PMCID: PMC8220552.

72. Huang L, Chen W, Tan Z, Huang Y, Gu X, Liu L, Zhang H, Shi Y, Ding J, Zheng C, Guo Z, Yu B. Mrc1(+) macrophage-derived IGF1 mitigates crystal nephropathy by promoting renal tubule cell proliferation via the AKT/Rb signaling pathway. Theranostics. 2024;14(4):1764–80. Epub 20240217. doi: 10.7150/thno.89174. PubMed PMID: 38389846; PMCID: PMC10879870.

73. Rodell CB, Arlauckas SP, Cuccarese MF, Garris CS, Li R, Ahmed MS, Kohler RH, Pittet MJ, Weissleder R. TLR7/8-agonist-loaded nanoparticles promote the polarization of tumour-associated macrophages to enhance cancer immunotherapy. Nat Biomed Eng. 2018;2(8):578–88. Epub 20180521. doi: 10.1038/s41551-018-0236-8. PubMed PMID: 31015631; PMCID: PMC6192054.

74. Wu N, Morsey BM, Emanuel KM, Fox HS. Sequence-specific extracellular microRNAs activate TLR7 and induce cytokine secretion and leukocyte migration. Mol Cell Biochem. 2021;476(11):4139–51. Epub 20210727. doi: 10.1007/s11010-021-04220-3. PubMed PMID: 34313894.

75. Tanaka R, Murakami Y, Ellis D, Seita J, Yinga W, Kakuta S, Kumano K, Fukui R, Miyake K. TLR7 responses in glomerular macrophages accelerate the progression of glomerulonephritis in NZBWF1 mice. Int Immunol. 2025;37(6):339–53. Epub 20250127. doi: 10.1093/intimm/dxaf005. PubMed PMID: 40401698.

76. Herzog S, Fragkou PC, Arneth BM, Mkhlof S, Skevaki C. Myeloid CD169/Siglec1: An immunoregulatory biomarker in viral disease. Front Med (Lausanne). 2022;9:979373. Epub 20220923. doi: 10.3389/fmed.2022.979373. PubMed PMID: 36213653; PMCID: PMC9540380.

77. Sprent J. Antigen-presenting cells. Professionals and amateurs. Curr Biol. 1995;5(10):1095-7. doi: 10.1016/s0960-9822(95)00219-3. PubMed PMID: 8548275.

78. Kaiser Y, Garris CS, Marinari E, Kim HS, Oh J, Pedard M, Halabi EA, Choi M, Parvanian S, Kohler R, Migliorini D, Weissleder R. Targeting immunosuppressive myeloid cells via implant-mediated slow release of small molecules to prevent glioblastoma recurrence. Nat Biomed Eng. 2025. Epub 20251022. doi: 10.1038/s41551-025-01533-2. PubMed PMID: 41125869.

79. Nie S, Song Y, Hu K, Zu W, Zhang F, Chen L, Ma Q, Zhou Z, Jiao S. CXCL10 and IL15 co-expressing chimeric antigen receptor T cells enhance anti-tumor effects in gastric cancer by increasing cytotoxic effector cell accumulation and survival. Oncoimmunology. 2024;13(1):2358590. Epub 20240524. doi: 10.1080/2162402X.2024.2358590. PubMed PMID: 38812569; PMCID: PMC11135867.

80. Agliardi G, Liuzzi AR, Hotblack A, De Feo D, Núñez N, Stowe CL, Friebel E, Nannini F, Rindlisbacher L, Roberts TA, Ramasawmy R, Williams IP, Siow BM, Lythgoe MF, Kalber TL, Quezada SA, Pule MA, Tugues S, Straathof K, Becher B. Intratumoral IL-12 delivery empowers CAR-T cell immunotherapy in a pre-clinical model of glioblastoma. Nature Communications. 2021;12(1):444. doi: 10.1038/s41467-020-20599-x.

81. Baker GJ, Chockley P, Zamler D, Castro MG, Lowenstein PR. Natural killer cells require monocytic Gr-1(+)/CD11b(+) myeloid cells to eradicate orthotopically engrafted glioma cells. Oncoimmunology. 2016;5(6):e1163461. Epub 20160316. doi: 10.1080/2162402x.2016.1163461. PubMed PMID: 27471637; PMCID: PMC4938363.

82. Tong Z, Kim C, Ross JL, Arvanitis C. Enhancing immune cell trafficking to brain tumors: Recent advances and therapeutic strategies. Neuro Oncol. 2026. doi: 10.1093/neuonc/noag075.

83. Luxenhofer R, Schulz A, Roques C, Li S, Bronich TK, Batrakova EV, Jordan R, Kabanov AV. Doubly amphiphilic poly(2-oxazoline)s as high-capacity delivery systems for hydrophobic drugs. Biomaterials. 2010;31(18):4972–9. Epub 2010/03/30. doi: 10.1016/j.biomaterials.2010.02.057. PubMed PMID: 20346493; PMCID: PMC2884201.

84. Sahn M, Weber C, Schubert US. Poly(2-oxazoline)-Containing Triblock Copolymers: Synthesis and Applications. Polymer Reviews. 2019;59(2):240–79. doi: 10.1080/15583724.2018.1496930.

85. Cho S, Rasoulianboroujeni M, Kang RH, Kwon GS. From Conventional to Next-Generation Strategies: Recent Advances in Polymeric Micelle Preparation for Drug Delivery. Pharmaceutics. 2025;17(10):1360. PubMed PMID: doi:10.3390/pharmaceutics17101360.

86. De la Cruz G, Nikolaishvili-Feinberg N, Gershon TR. Automated Immunofluorescence Staining for Analysis of DNA Damage and Apoptosis in Brain Sections. Methods Mol Biol. 2023;2583:55–61. doi: 10.1007/978-1-0716-2752-5_6. PubMed PMID: 36418725.

