## Supplementary figures and images for "Myeloid compartment reprogramming through nanoparticle-delivered resiquimod blocks paracrine growth support and activates phagocytosis to slow tumor progression in endogenous mouse medulloblastoma and diffuse midline glioma models"

### Supplementary Figure 1

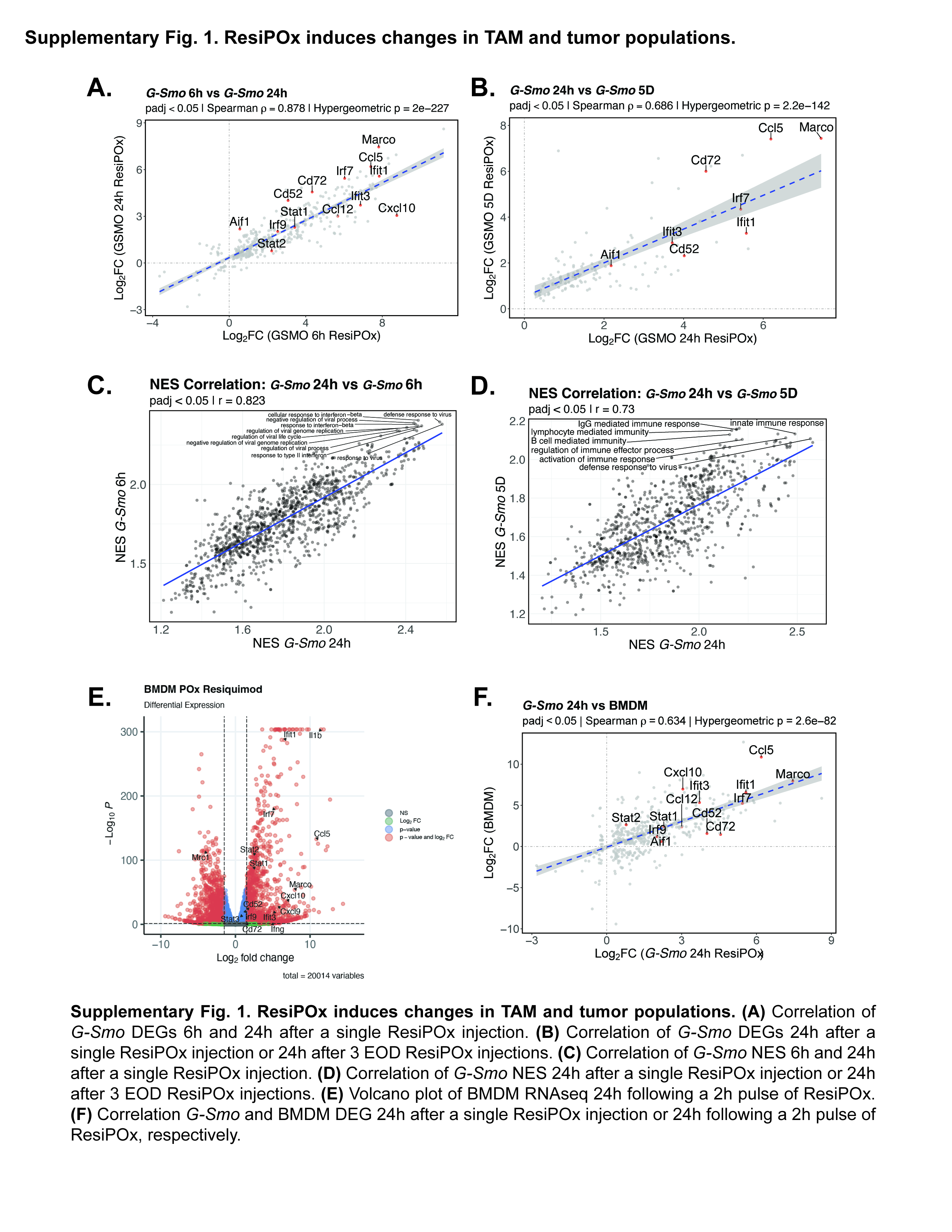

### Supplementary Figure 2

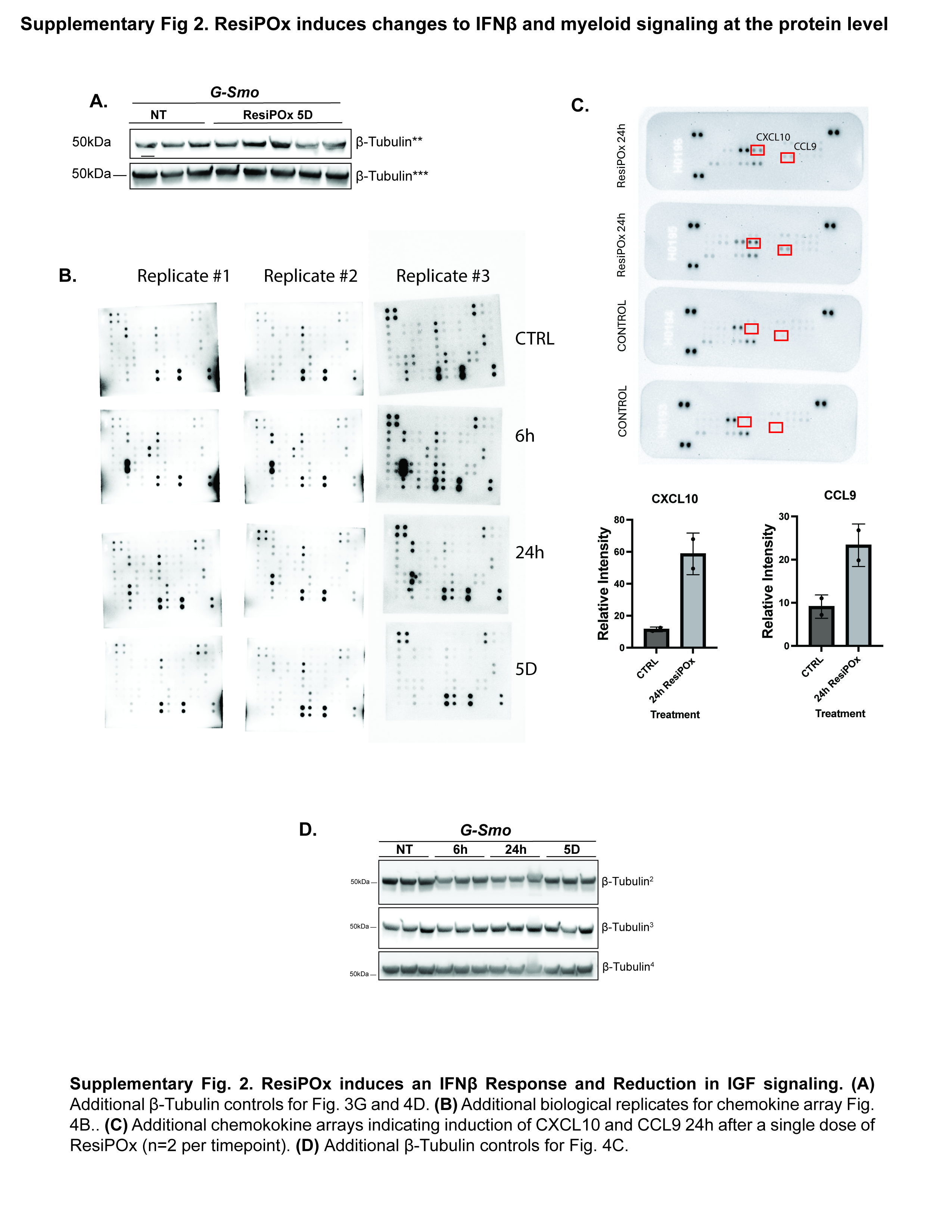

### Supplementary Figure 3

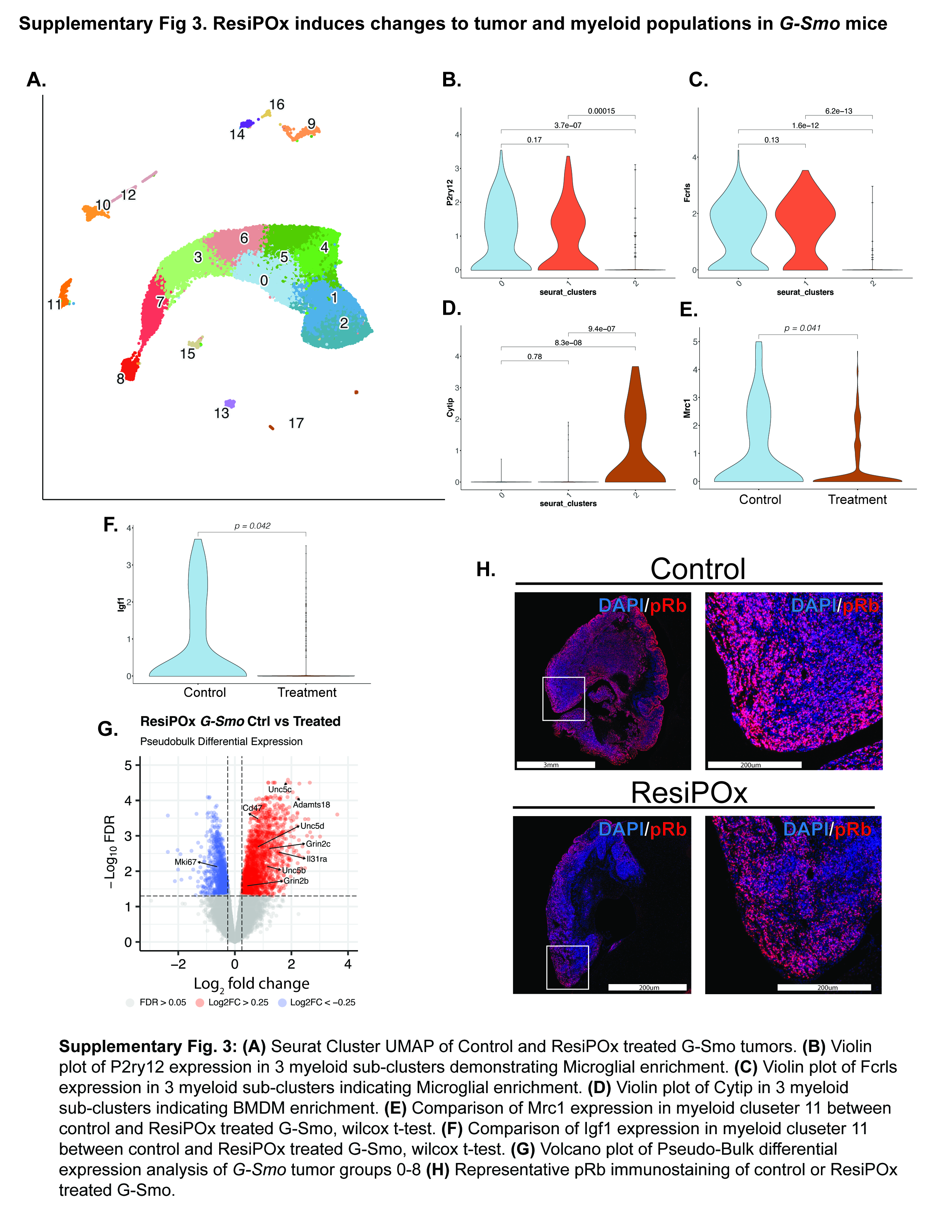

### Supplementary Figure 4

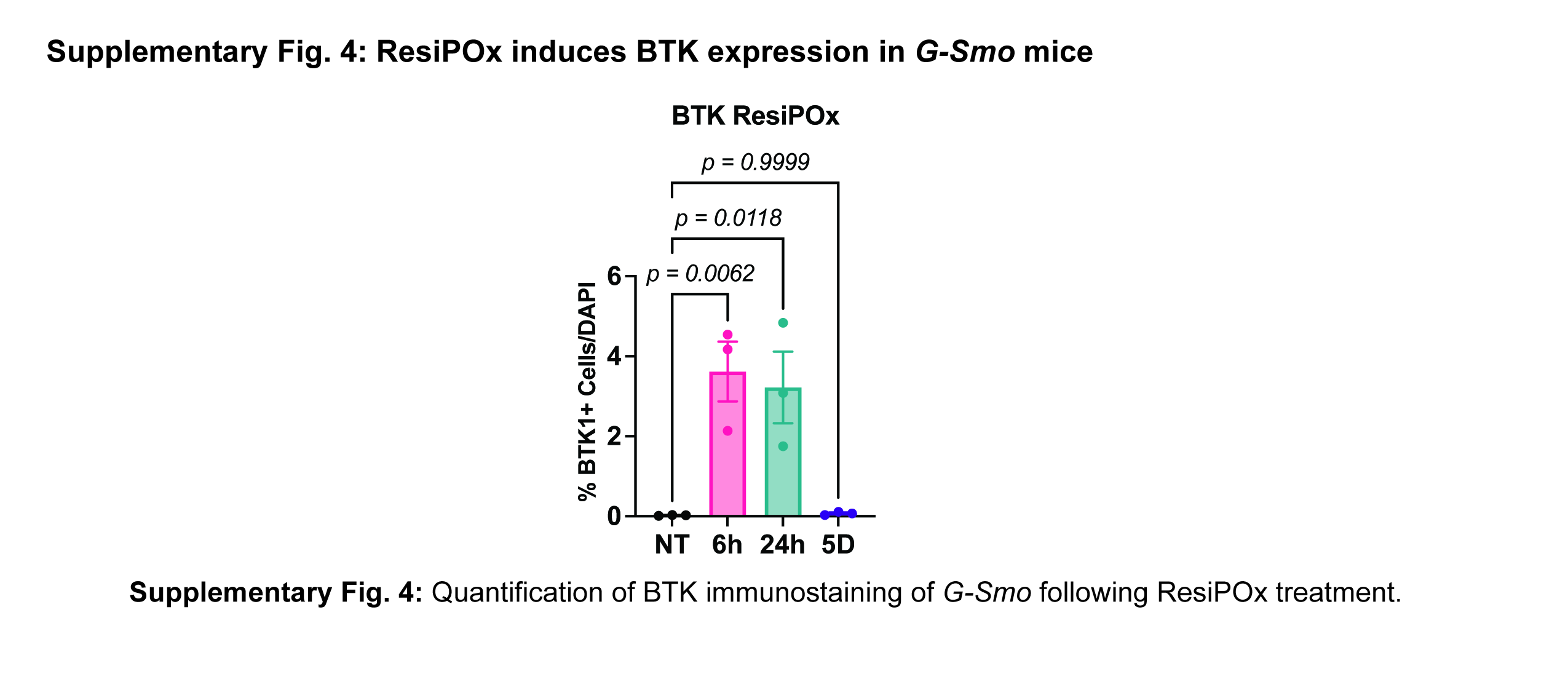

### Supplementary Figure 5

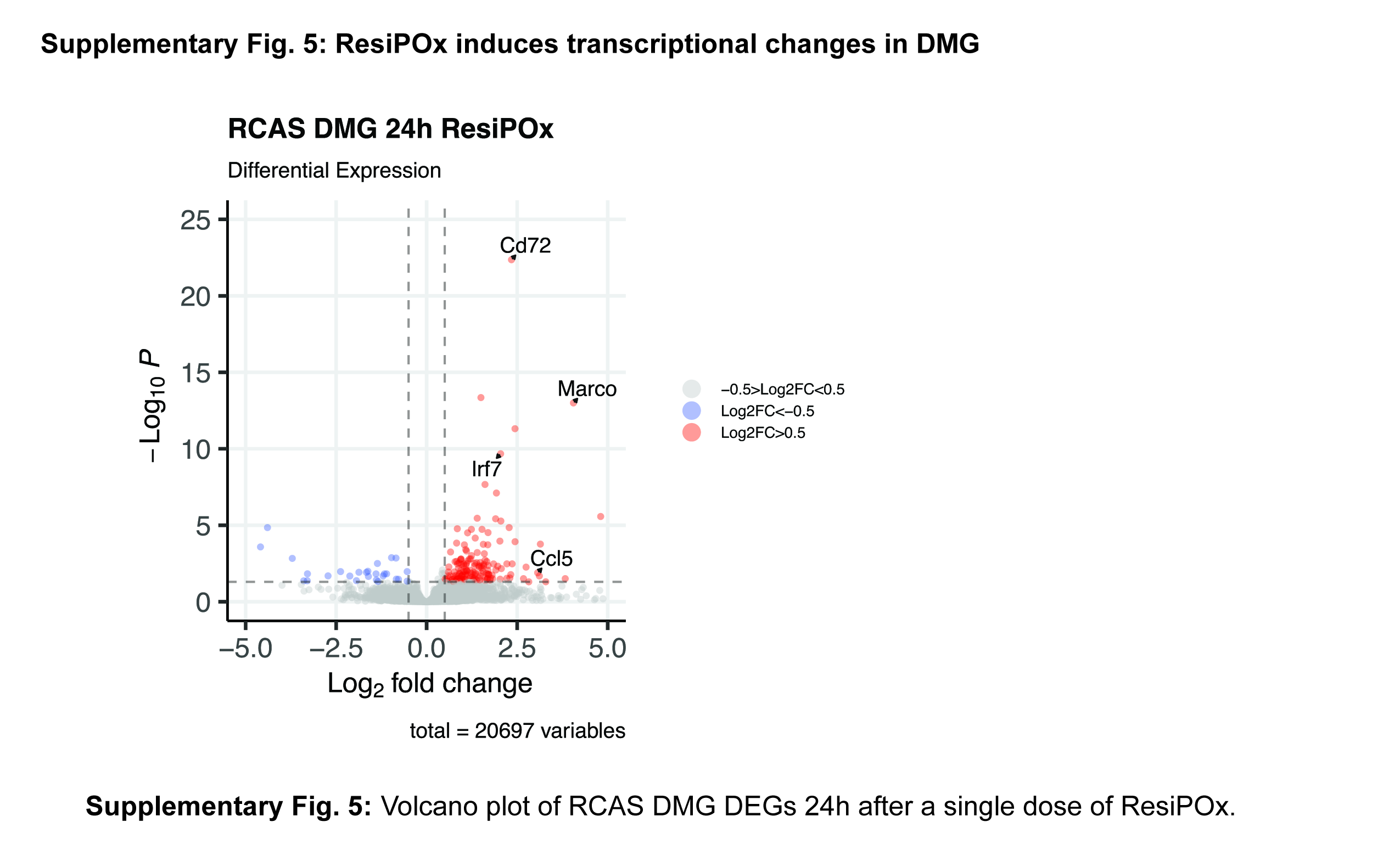

### Supplementary Figure 6

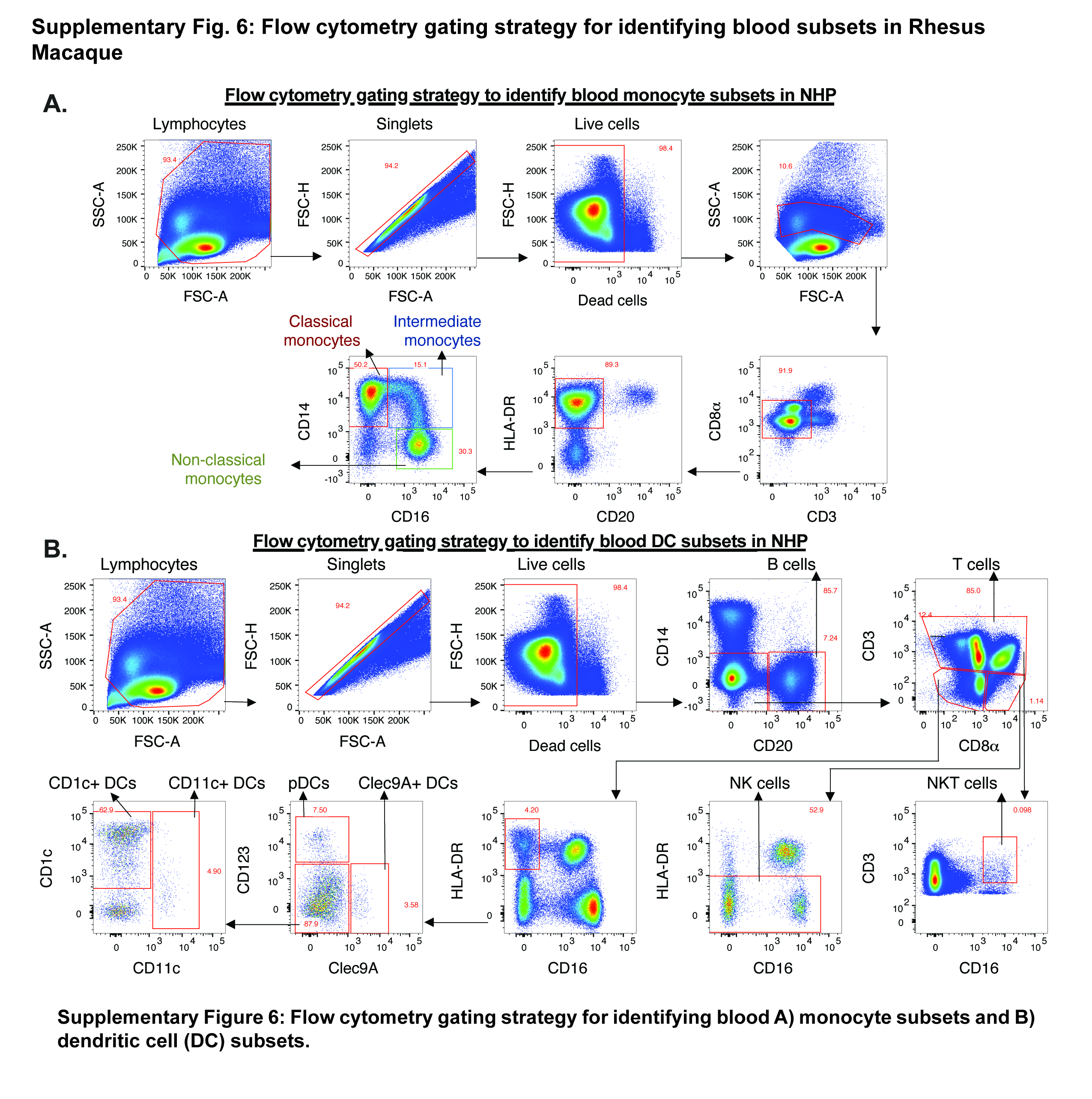

### Supplementary Figure 7

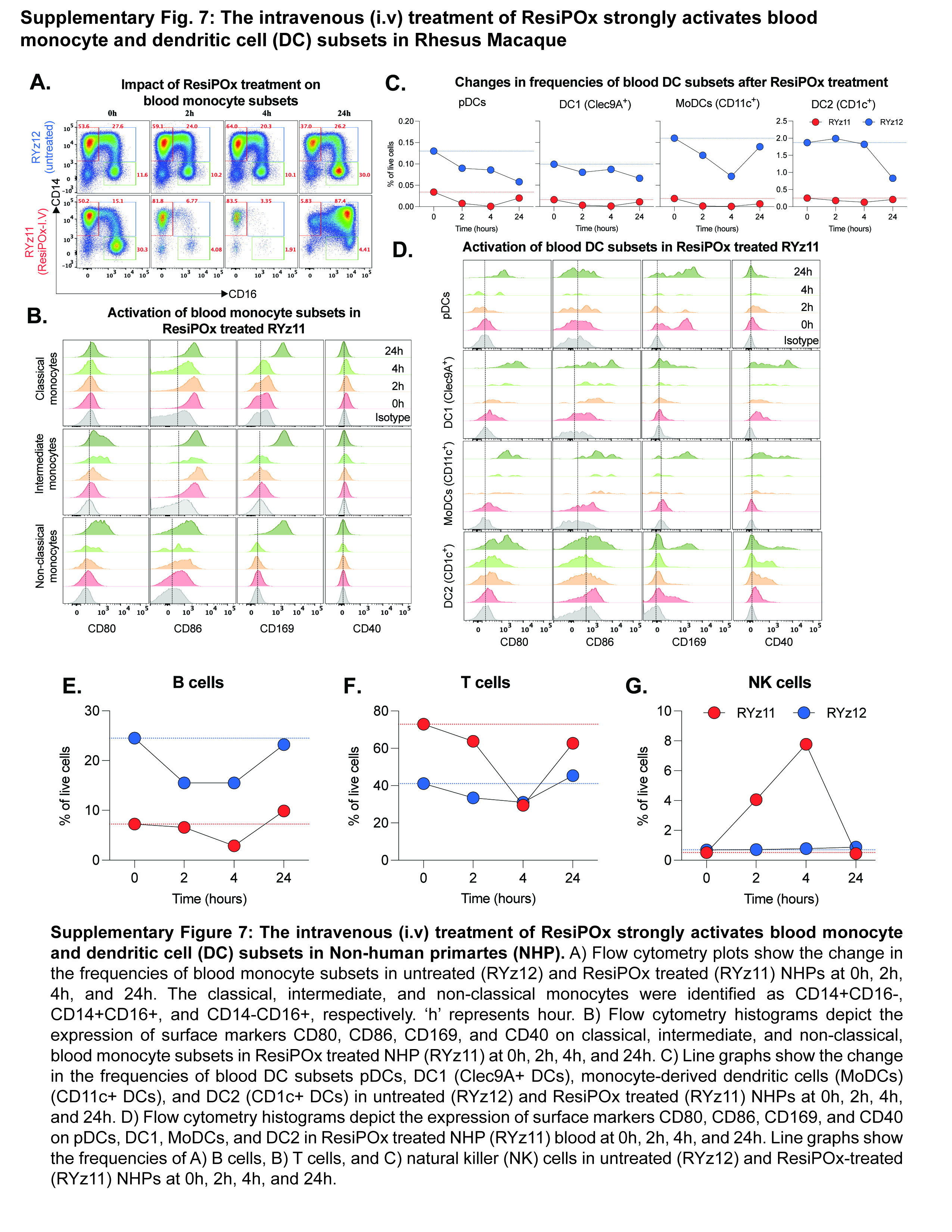

### Supplementary Figure 8

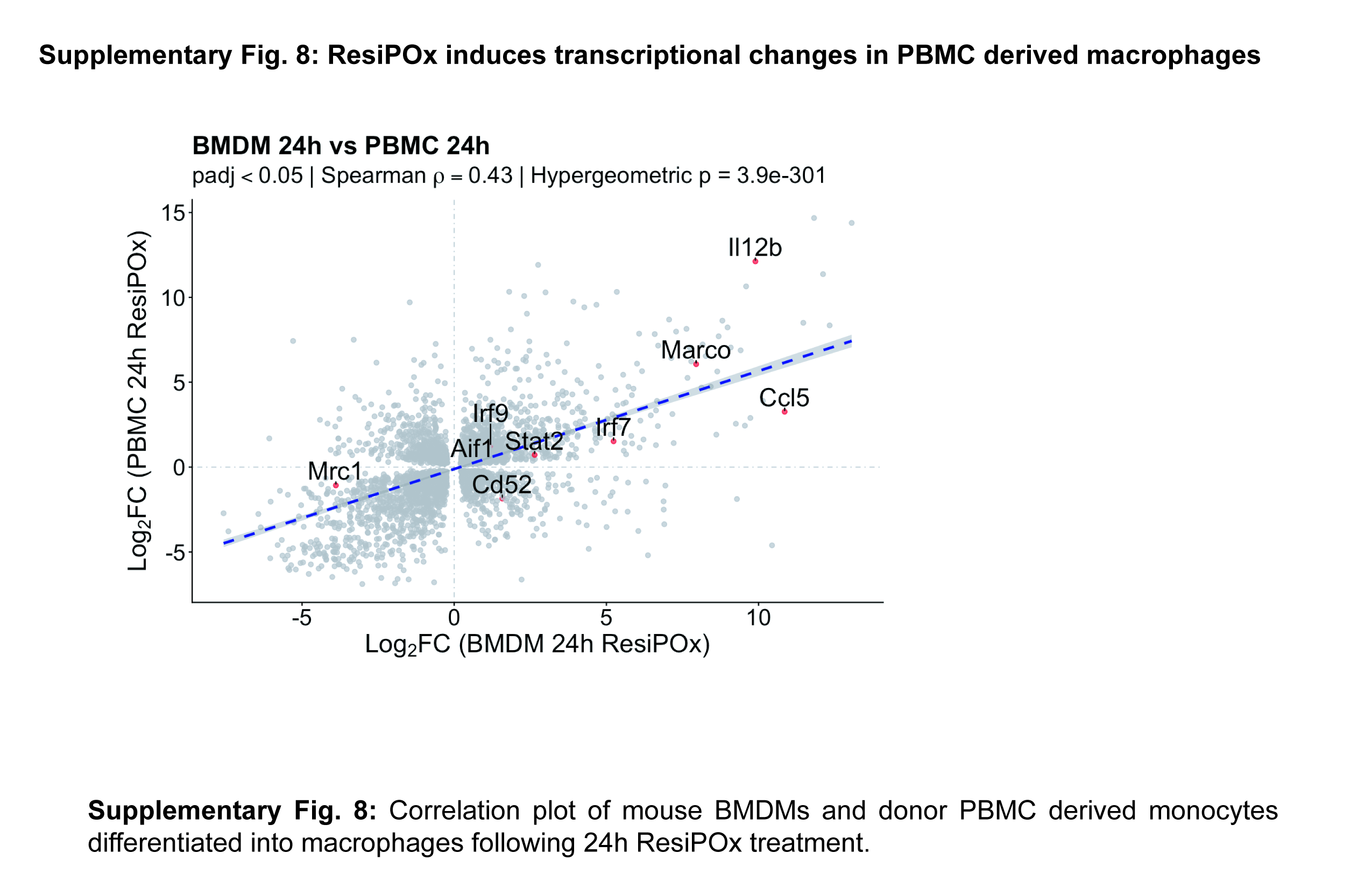
